# Mass-specific single molecule pull-down from complex mixtures with bilayer-assisted mass photometry

**DOI:** 10.64898/2026.02.07.704484

**Authors:** Manish S Kushwah, Raman van Wee, Jan Christoph Thiele, Eric D. B. Foley, Justin L.P. Benesch, Philipp Kukura

## Abstract

Determining the identity, quantity and interactions of proteins is fundamental to studies of biomolecular function and (dis)regulation in disease. Here, we combine mass photometry with supported lipid bilayers (SLBs) to specifically recruit proteins of interest from complex mixtures such as cell lysate and demonstrate sensitive detection and mass measurement of single biomolecules and their complexes without prior purification. Extending to heterologously expressed proteins, we provide rapid information on oligomerisation and concentration in lysate. Introduction of polyethylene glycol in the SLBs passivates against a variety of cell lysates and serum, allows specific recruitment of proteins from complex mixtures and mass-specific detection and affinity measurement at sub-nM concentrations. Our approach preserves the mass resolution and accuracy of mass photometry, thereby enabling visualisation and quantification of oligomeric states, interactions and complexes directly from cell lysates.

## Introduction

The majority of biophysical and bioanalytical studies rely on protein purification and quantification. Most commonly, this is achieved by affinity-based purification followed by absorbance measurements to determine protein concentration, SDS-PAGE gel electrophoresis, and Western blotting to confirm protein identity. While very well established, this workflow remains laborious and especially for Western blotting, challenging. Even if all steps are performed correctly, they provide little information on the state of the protein, because SDS-PAGE and Western blotting are performed in denaturing conditions, where non-covalent interactions have been disrupted.

In an ideal scenario, one would be able to study proteins and the complexes they form directly from complex mixtures in a rapid and specific fashion. Single-molecule pull-down (SiMPull) has emerged as a powerful alternative to rapidly detect individual protein complexes, quantify their abundance and kinetics, while using minimal sample volumes down to a few cells^1–4^. In a standard SimPull experiment, an antibody against a protein of interest is immobilised on a passivated surface after which cell extract is added and subsequently washed away. Protein detection takes place either directly *via* a genetically expressed fluorescent tag, or by flushing in a fluorescently labelled antibody against a different epitope^2,5^. Since its inception^1^, SimPull has been optimized extensively, making it now possible to pull-down proteins using 3 orders of magnitude less cell lysate than needed for bulk assays^3^, study DNA-protein interactions^6^ and pull-down proteins from single cells^4,7,8^.

Fluorescence-based detection at the single-molecule level in SiMPull, however, either requires fluorescent labelling, or remains an indirect readout, because it reports on the presence of a second antibody on the surface^9^. This makes the measurement susceptible to non-specific interactions, complicating both detection and quantification. In addition, it struggles to quantify oligomerisation and suffers from practical limitations to simultaneously image multiple fluorophores needed to quantify the stoichiometry of multi-protein complexes^10–12^. For small oligomers, stepwise photo-bleaching can be used to detect oligomerisation^13^, however, reproducibility and ease of experiment remains challenging. Moreover, extensive functional characterisation of proteins labelled with genetically encoded fluorescent proteins further prolongs and complicates experimental process.

Recently, scattering-based approaches have been developed to mitigate the challenges posed by fluorescence-based SiMPull^14,15^. These approaches are capable of label-free detection of single proteins from cell lysate; however, detection of oligomerisation and stoichiometry of protein complexes remains out of reach, and the binary read-out means specificity relies entirely on the performance of the passivation and functionalisation strategy. Mass photometry (MP)^16–18^, is in principle a viable alternative to overcome these limitations. In a traditional MP assay, however, label-free detection and mass measurement takes places after non-specific binding from solution onto a glass surface (Fig. 1a), These binding events cause a change in local refractive index and thus reflectivity of the glass-water interface, which are quantified (Fig. 1a). In purified samples, different oligomers or complexes can be distinguished in the resulting mass distributions, representing the oligomeric composition of the purified protein (Fig. 1b).

**Figure 1.**
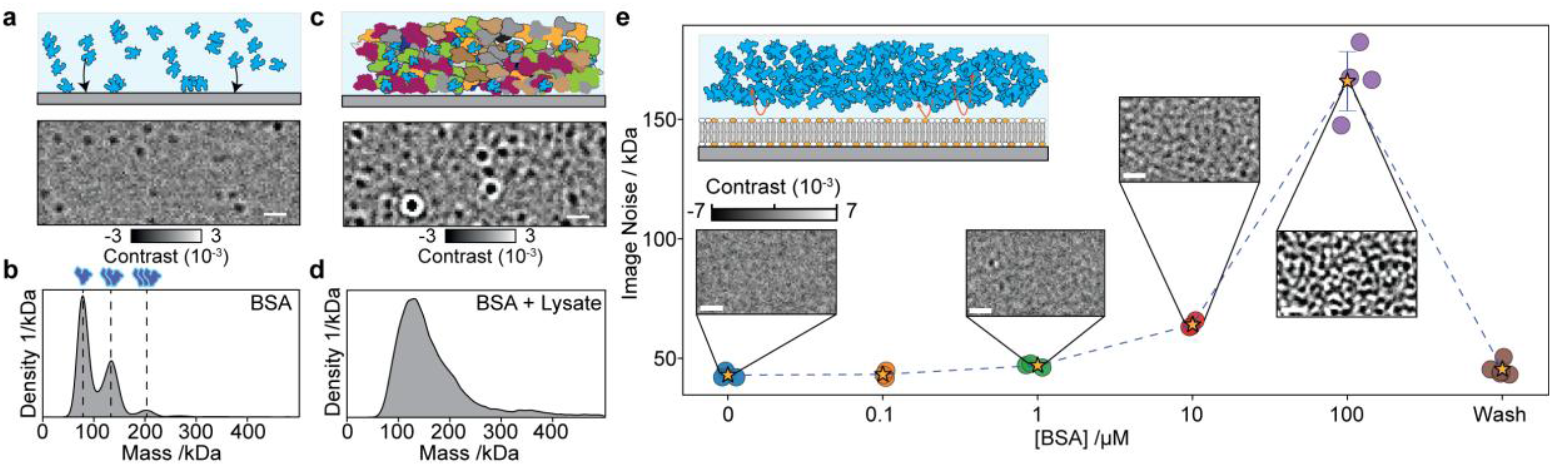
Glass passivation against non-specific binding in Mass Photometry using supported lipid bilayers (SLBs). a) Standard MP experiment and ratiometric MP image of BSA (25 nM) on microscope cover glass. b) Resulting MP histogram (dashed lines indicate BSA oligomers. c) Equivalent experiment in the presence of *E. coli* lysate. d) Corresponding MP distribution. e) Bilayer noise quantification where increasing BSA concentration is added to the glass supported bilayer. The star indicates the mean noise from multiple datasets.

Performing the same experiment in the presence of the contents of even a relatively simple cell such as *E*.*coli* lysate, which contains about 1000-3000 types of proteins^19^, is essentially impossible (Fig. 1c). The non-specific nature of the glass-protein interaction causes all proteins to bind to the glass surface making it impossible to distinguish species of interest in a complex mixture (Fig. 1c). The result is a broad, unresolved distribution representative of the relative abundances of biomolecules and their masses in the lysate (Fig. 1d), meaning all specific and quantitative information normally available to MP is lost.

In principle, these problems could be overcome by PEG passivation of the glass surface and performing an inverse SiMPull strategy, where unbinding of specifically pulled down species is detected after a washing step^20^. Such a strategy, however, requires specific detachment of bound molecules, and relies on high differential passivation to ensure that unbinding events are indeed specific. In this work, we demonstrate the advantages of supported lipid bilayers (SLBs) combined with the development of a robust PEG-based bilayer passivation method for MP-based recruitment and characterisation of proteins, stoichiometry of their complexes and interactions directly from complex mixtures such as mammalian cell lysates without prior purification steps.

## Results

### Bilayer-mediated passivation and specific detection with mass photometry

SLBs provide a densely-packed, two-dimensional array of lipids on top of a glass surface, which can be simultaneously used for passivation^21^ and specific recruitment^22–25^. SLBs support lipid diffusion^26–29^, which allows for dynamic detection of species, enabling the use of mobility as an additional differentiating factor between specifically and non-specifically bound species. SLBs have not been used in the context of fluorescent SiMPull, because detection of rapidly diffusing species is challenging in the context of a finite photon-budget^12,30^. These limitations do not apply to MP^26,31^, where bilayer integrity and pulled-down biomolecules can be detected simultaneously and label-free detection overcomes the trade-off between spatial and temporal resolution.

To explore the performance of SLB passivation in an MP assay, we tested non-specific binding of bovine serum albumin (BSA) to an SLB containing 89 mol% DOPC, 10 mol% DOPS, and 1 mol% DGS-NTA (PS-NTA bilayers). We chose BSA as a first test of SLB passivation capacity as BSA non-specifically binds to variety of surfaces and is routinely used for surface passivation in single molecule assays, western blots and ELISA^32–35^. To assess the effect of BSA binding on bilayers, we incubated SLBs with increasing BSA concentrations (0-100 µM) and quantified the image noise expressed as protein mass equivalent as a function of BSA concentration (Fig. 1g). The SLBs successfully prevented BSA binding up to 100 nM. At 1 µM solution concentration, first signatures of diffusing proteins appeared and overwhelmed the MP images at 100 µM (Fig. 1g; Supplementary Videos 1-2). In both cases, we found that at 100 µM BSA SLBs are unusable for MP (Fig. S2, S3), however, after a simple wash corresponding to an effective BSA concentration 2 pM, the image noise returned to the baseline value and we could not detect any signatures of residual proteins bound to the SLB (Fig. 1g; Supplementary Video 3). These results indicate that SLBs can sustain prolonged exposure to non-specific proteins that can be subsequently removed by washing.

### Bilayer-mediated single molecule pulldown from lysates and protein quantification

Removal of non-specific binding on SLBs suggests that proteins can be specifically recruited with functionalised lipids, even in the presence of substantial amounts of background species. To test this possibility, we mixed 1 nM purified 6xHis Dyn1ΔPRD (Dyn1, Fig. 2a), which shows an oligomeric distribution in solution (Fig. 2b) with 100 µM BSA and incubated the mixture on an SLB with the same lipid composition as above for 5 minutes (Fig. 2c). After washing with assay buffer, we found bound species, whose increasing mass and decreasing mobility matched that of Dyn1 dimers, tetramers, and hexamers (Fig. 2d)^26^. The readout outlines a key advantage of simultaneous mass and mobility measurement: in contrast to non-specific background, pulled down targets exhibit distinctive molecular mass and diffusion, determined by the number of lipid contacts to the SLB (Fig. 2d; Supplementary Video 4).

**Figure 2.**
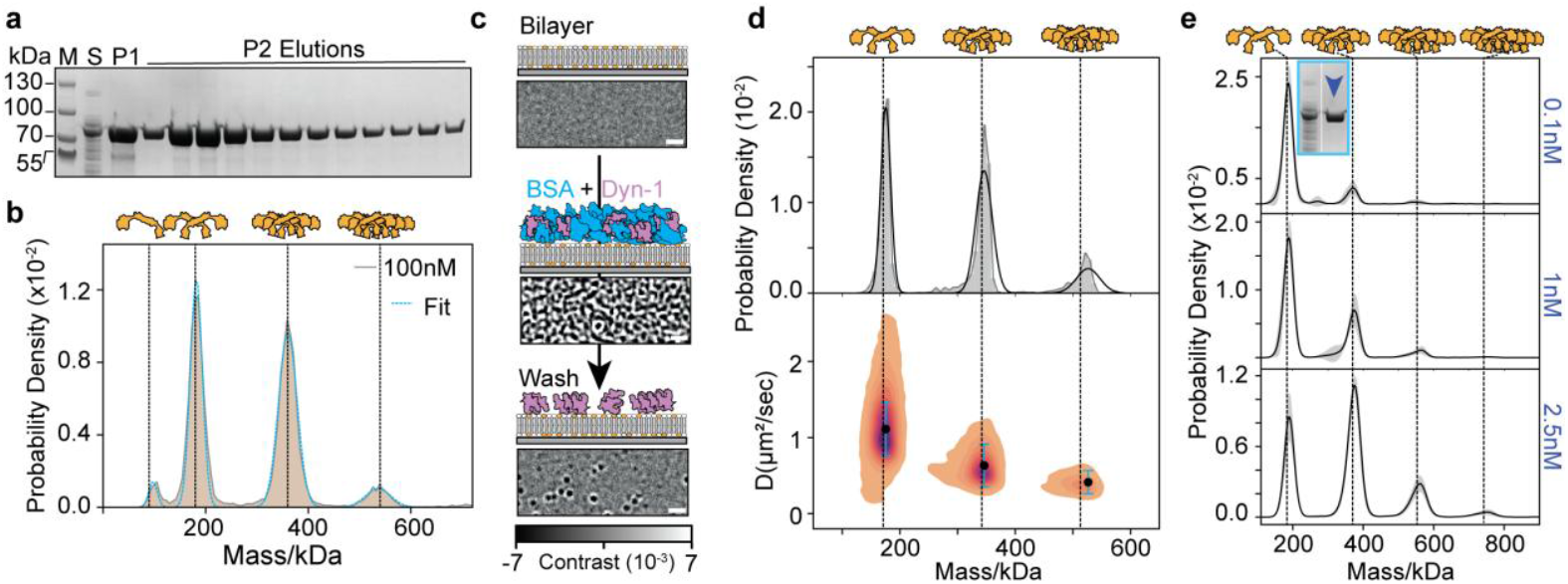
SLB-mediated single-molecule pulldown. a) SDS-PAGE gel showing purified Dyn1 (P1, IMAC elution, P2 Strep-tag elution) and clarified E.coli lysate (S) containing over-expressed Dyn1. b) Pooled P2 at 100 nM monomer concentration. Vertical lines indicate expected mass of Dyn1 oligomers. c) Schematic and representative background subtracted images of a clean SLB (top), after addition of Dyn1 (1 nM) in the presence of 100 µM BSA (middle) and following a buffer wash after 5 min incubation (bottom). Scale bar = 1 µm. d) Corresponding mass-diffusion plot of BSA-spiked Dyn1. e) Mass distributions as a function of Dyn1 concentration in the presence of BSA. For each concentration, data from 3 movies at 3 different SLBs was used. Black lines and grey areas indicate the mean and standard deviations.

Repeating this experiment at increasing Dyn1 concentrations mixed with 100 µM BSA, followed by washing revealed a steady increase of larger oligomers with increasing analyte concentration, demonstrating that our assay is not only specific, but also maintains intermolecular interactions observed in solution (Fig. 2e). These results suggest that SLB-based capture efficiently isolates biomolecules of interest from a large background and does so in a fashion amenable to MP-based analysis.

To test to which degree this approach remains viable in the presence of a more complex cellular lysate background, we added increasing concentrations of Dyn1-containing *E. coli* lysate (Fig. 3a) to our SLBs, resulting in a steady increase in detected particles. In this case, due to the high expression levels of Dyn1, no washing steps were required to reveal the analyte of interest. The resulting oligomeric distributions closely matched those observed for purified Dyn1 (Fig. 3b, compared to Fig. 2e). We could further quantify the oligomeric distribution of Dyn1 from lysate, which evolved with increasing lysate concentration in a similar manner as purified Dyn1 and remained stable after washing away lysate (Fig. 3b, SI Fig 4-5, 7) showing the importance of the assay in studying protein interactions straight from lysates. This oligomeric distribution is consistent with previous data from cells^36^ and in dynamic MP-based in-vitro assays^26^. The absence of any mass beyond the oligomers of Dyn1 further suggests that other *E. coli* lysate proteins, such as FtsZ or bacterial Dynamins, do not interact strongly enough with the lipid surface of this composition at the lysate concentrations tested to create a detectable MP signal.

**Figure 3.**
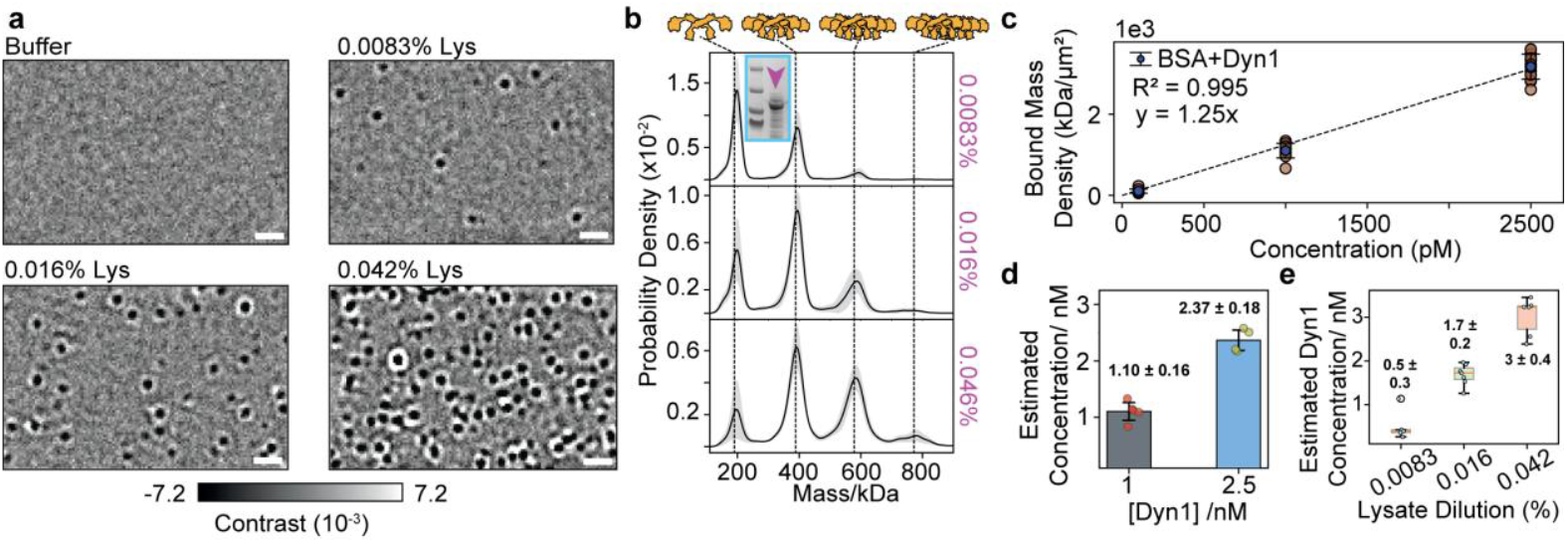
MP of overexpressed proteins directly from cell lysate. a) Representative MP images of SLBs with increasing E.coli lysate concentration. Scale bar: 1 µm. b) Corresponding mass distributions at increasing lysate concentrations. c) Bound mass density as a function of Dyn1 concentration in the presence of 100 µM BSA. d) Dyn1 concentration measurement for Dyn1-spiked lysates using the calibration from (c). e) Equivalent for data in a,b.

**Figure 4.**
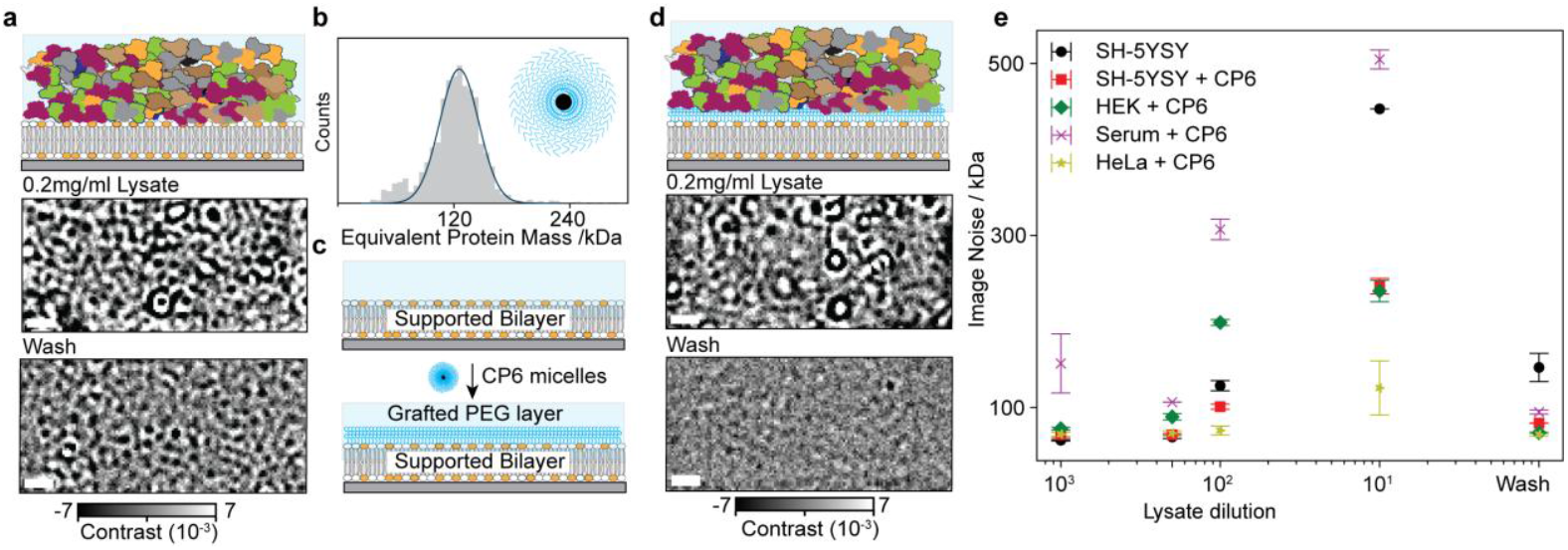
Cholesterol-PEG (CP6)-600 micelle-assisted PEG insertion and improved SLB passivation to cell-lysates. a) Concentrated ShSy5y lysate addition to bilayers results in non-specific binding that cannot be remove by washing. Scale bar: 1 µm. b) Mass histogram of Cholesterol - PEG600 (CP6) at 10 µM. c) Schematic of CP6 micelle-mediated PEG insertion into preformed SLBs. d) CP6-PEG passivation against lysate. e) Efficacy of CP6-mediated bilayer passivation against various cell lysates and bovine foetal serum.

The dependence of Dyn1 oligomerisation on solution concentration combined with the similarity between the measurement in lysate with purified protein suggests that it is possible to determine the Dyn1 concentration in *E. coli* lysate directly. To test this, we evaluated the repeatability of quantifying the total mass of proteins bound to the SLB. We used the oligomeric distribution from the Dyn1+BSA experiment (Fig. 2e) and extracted the bound mass density on SLBs. We then integrated the abundance of each oligomeric species weighted by its stoichiometry (number of monomers), summed across all species, and normalized to the total acquisition time and the field-of-view to obtain a measure of total bound protein.

Next, to obtain a calibration between bound mass density and Dyn1 solution concentration, we varied the concentration of Dyn1 in the presence of 100 µM BSA, incubated each concentration on a PS-NTA SLB for 5 minutes, and washed unbound Dyn1 and BSA. We found that the total bound mass increased linearly with Dyn1 concentration in the Dyn1+BSA mixture reproducibly yielding 1243 ± 100 kDa / μm^2^ / nM Dyn1 (best fit slope ± standard error of the fit, Fig. 3c). We then used the resulting concentration to bound mass calibration to determine the concentration of 1 and 2.5 nM Dyn1 spiked into 100 μM BSA, showing good agreement (Fig. 3d). This calibration also predicted Dyn1 concentration with good accuracy in a separate experiment where pure Dyn1 was added to SLBs (SI. Fig 8), showing that background species do not significantly inhibit Dyn1 recruitment to SLBs. Using this calibration on our lysate dilution showed the expected linear trend (Fig. 3e) and yielding 8 ± 1 µM (μ ± σ) Dyn1 concentration.

### PEG-based passivation for single-molecule pulldown from mammalian lysates

For overexpressed proteins, high dilution levels can be used while maintaining sufficient statistics for protein detection and quantification. For native expression levels, however, such high levels of dilution render most proteins undetectable, requiring the use of low or undiluted complex mixtures. Now, membrane-active proteins can bind to and/or change bilayer topology, rendering MP measurements on SLBs challenging. Indeed, we found substantial residual binding after adding SHSy5y cell lysate at a total protein concentration of ∼0.2 mg/ml even after multiple washes (Fig. 4a), making further characterisation by MP impossible and calling for additional passivation.

Previous approaches have used PEG-modified lipids to improve bilayer passivation^37–40^, but this did not improve performance in our highly sensitive assays (Fig. S10), calling for even higher levels of passivation. Drawing inspiration from membrane-targeting of soluble proteins via covalent modifications with a hydrophobic moiety^41^, we delivered PEG to pre-formed SLBs surface via cholesterol, rather than incorporate it directly through modified lipids. We chose cholesterol-PEG600 (CP6), which has previously been shown to bring cholesterol to cells^42–44^ and to increase the solubility and lifetime of lipid nanoparticles for drug delivery^45,46^. We hypothesized that akin to detergent monomer incorporation into a bilayer via its hydrophobic tail^47^, CP6 would deliver PEG to bilayers with a density controlled by the solution concentration (Fig. 3c; Fig. S10). In contrast to cholesterol which is sparingly soluble (2×10^-5^ mg/ml)^48^, the amphipathic nature of CP6 is amenable to micelle formation and increases its solubility up to 60 mg/ml. An MP landing assay of CP6 at 10 µM showed that CP6 produced a monodisperse distribution of micelles of an equivalent protein mass of 126 kDa, corresponding to ∼126 lipids per micelle (Mw CP6 = 1 kDa, Fig. 4b), comparable to reported values for similar molecules^49^. CP6 passivation did not affect Dyn1 pulldown from *E*.*coli* lysate on PS-NTA SLBs (SI. Fig 12).

Our PS-NTA bilayers (Fig. 4c) incubated for 5 minutes with 20 µM CP6 showed improved passivation after incubation with cell lysate at 0.2 mg/ml after washing (Fig. 4d). To test the robustness of CP6-mediated bilayer passivation, we exposed CP6-fortified SLBs to a variety of samples, including SHSy5y, HeLa, and HEK lysates and foetal bovine serum at increasing concentration before washing. The background noise of CP6-fortified bilayers returned to the baseline value for all samples, whereas SLBs without CP6 showed significantly higher noise (Fig. 4e; Fig. S11). These results show that addition of CP6 offers additional and robust passivation of SLBs against cell lysates and serum, bringing quantification of proteins at native expression levels within reach.

Equipped with a well-passivated SLB, we investigated the performance of our assay in the context of a more traditional coIP/SimPull assay, but one where we can control the exact amount of target analyte. For this, we designed an antigen-antibody system in which the Fc domain of a primary antibody interacts with the Fab domain of secondary antibodies (SecAbs) used to visualise specific proteins in western blots or ELISA assays^50^. We chose a 6xHis-tagged Fc dimer (hFc; mol. Wt. 28 kDa) of a primary Human IgG1 with two secondary antibodies (SecAbs) specific to human IgG1 Fc domain (αhFc-IgG1 (hIgG1) & αhFc-IgG2 (hIgG2), Fig. 5a, b). To study any species cross-reactivity, we also chose a secondary antibody specific to Fc dimer of mouse IgG1; anti-mouse Fc (mIgG).

**Figure 5.**
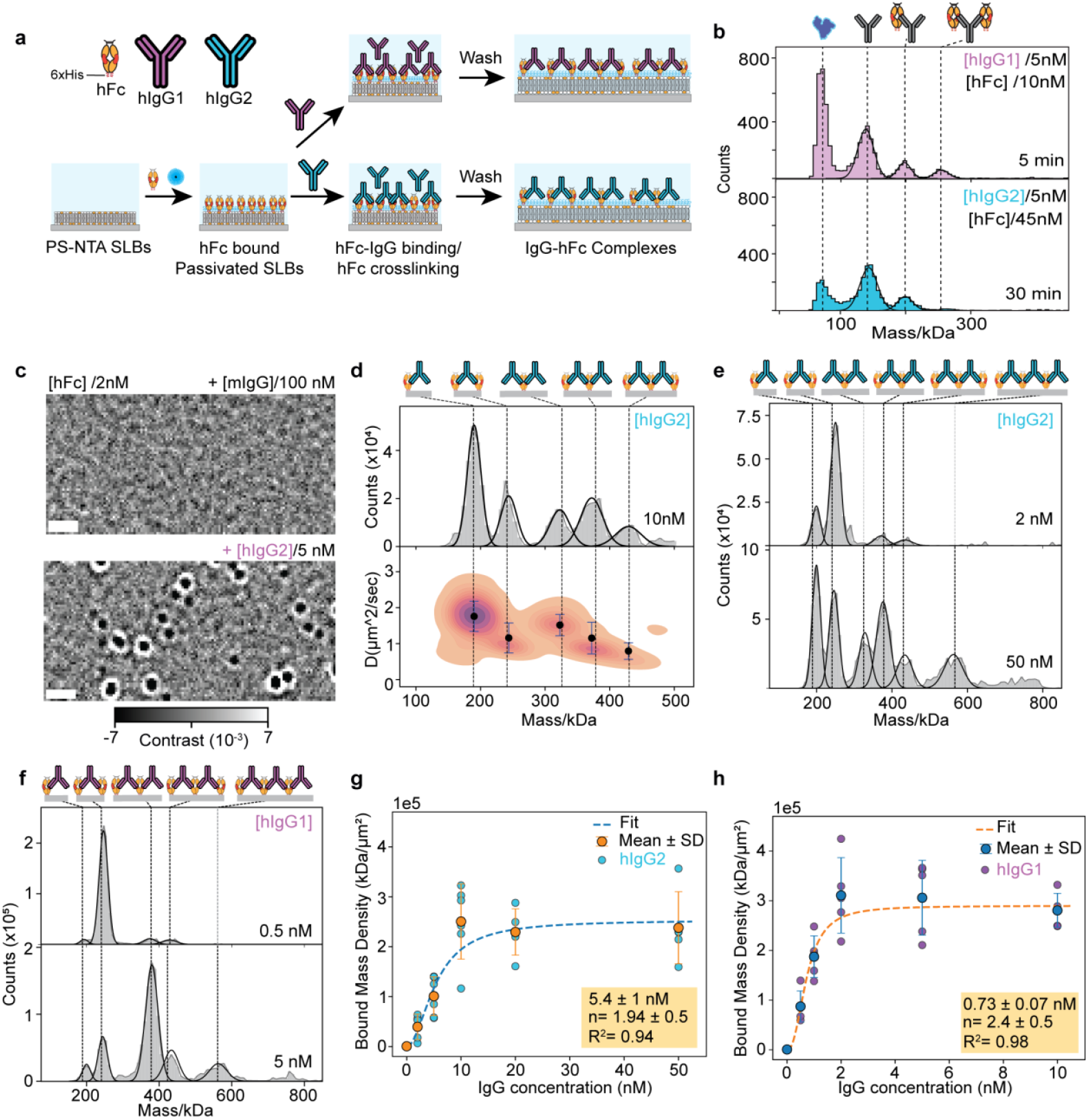
CP6-passivated SLBs provide an estimate of antigen-antibody affinity. a) Schematic of hFc mediated pulldown of antibodies on CP6 passivated PS-NTA bilayers. b) Landing assay showing the affinity of hIgG1 and hIgG2 to hFc in solution. Incubations times are reported in the corresponding panels (see also Fig. S13). c) Cross-reactivity/non-specific binding of mIgG to hFc (see also Fig. S14). Scale bar: 1 µm. d) Mass-diffusion plot for hIgG2 recruitment and complex formation under limiting hFc concentration. e-f) hIgG2 (e) and hIgG1 (f) concentration dependent changes in bound mass and hFc-hIgG complexes on CP6 SLBs containing limiting hFc concentration. g-h) Titration of hIgG2(g)/1(h) on CP6 passivated SLBs containing limiting hFc concentration to estimate the affinity for two antibodies. For each antibody, *K*_D_, hill coefficient, and R^2^ shown.

In a first step, we determined the mass distributions of the hFc and SecAbs. The hFc yielded a monodisperse distribution centered at 56 kDa, indicating that hFc forms a dimer at 5 nM (Fig. S13). Both SecAbs exhibited a molecular weight of ∼140 kDa (Fig. S13). Next, we determined the affinity of the two SecAbs to hFc using MP^51^ at 5 nM IgG. For hIgG1, we observed significant complex formation already at 10 nM hFc, indicative of a strong interaction (Fig. 5c). By quantifying the relative amounts of unbound, one-bound and two-bound hIgG1-hFc, we found an apparent affinity of *K*_D, app_ ≈ 2 x 10^1^ nM. For hIgG2, complex formation was less pronounced, even at 45 nM hFc, yielding a weaker affinity: *K*_D, app_ ≈ 3 x 10^2^ nM. Whereas the manufacturer reports sub-nanomolar EC50 values in an ELISA assay, these measurements are ensemble-average values of the multivalent interactions whose value could affected by avidity effects. With MP, we can resolve heterogeneous complexes directly in solution and quantify the underlying affinities.

To study the hFc-SecAbs interaction on SLBs, we first tested for non-specific binding and species cross-reactivity. We added 2 nM hFc to SLBs and without washing excess hFc, added increasing concentrations of mIgG. We found no evidence for antibody binding even at 100 nM non-specific mIgG, however, addition of 5 nM of specific antibody, hIgG2, led to substantial binding and oligomerisation (Fig. 5c, SI Fig. 14).

To study specific antibody binding, we recruited 2 nM hFc to CP6-fortified PS-NTA SLBs, washed excess hFc after 5-minute incubation and added specific SecAbs. We observed 1:1 and 2:1 IgG:hFc complexes and higher-order complexes containing multiple hFcs crosslinked by hIgG2 (Fig. 5d). As the mobility of species scales inversely with the membrane contact area^26,29^, the diffusion coefficient is consistent with the number of Fc dimers in a complex. Combined, the mass and mobility measurement enable us to identify all species in the distribution (Fig. 5d).

Since the total mass of species pulled down onto the SLB and the degree of hFc crosslinking by IgGs depends on the affinity of the antigen-antibody interaction, we reasoned that our platform in principle enables the quantification of this affinity directly from cell lysate. We thus compared the mass histograms at equal hIgG concentrations (10 nM) added to CP6 fortified SLBs pretreated with 2 nM hFc dimer and washed to remove excess hFc. hIgG1 formed much larger complexes than hIgG2, indicative of a higher affinity, as also observed in the landing assay (Fig. 5e-f; see Fig. S15 and S16 for hFc-IgG complexes, their molecular weight and respective stoichiometry). This comparison shows that the measured mass distribution indeed depends on IgG-Fc affinity.

To quantify the respective affinities for these two SecAbs, we next performed titrations for a fixed Fc dimer concentration and quantified the bound mass density of recruited species. Both for hIgG1 and hIgG2, the total mass on the bilayer increases with concentration and reaches a similar plateau value at ∼2.5 x 10^5^ kDa/μm^2^, likely limited by the number of capture complexes on the SLB (Fig. 5g-h). A fit of the data with the Hill-equation reveals an apparent affinity of 0.8 ± 0.2 nM for IgG1 and 5.4 ± 2.2 nM for IgG2 (both best fit ± standard error of the fit). These apparent affinities are substantially tighter than the corresponding solution affinities for both SecAbs, which we attribute to a combination of the effect of dimensionality reduction and enhanced cooperativity on a surface compared to free solution.

Given the difference in SLB-recruitment as a function of target affinity, our assay could in principle be used to detect antibody from lysates^52–54^ or determine the amount of IgG directly in cell lysate for a known affinity or vice versa, advantageous in antibody production and QA/QC^55^. To test this, we spiked SecAbs into HeLa lysate and incubated this mixture onto CP6-fortified PS-NTA SLBs with hFc (Fig. 6a). We found that regardless of concentration and affinity, hIgG1/2 complexes recruited from lysate on CP6-passivated SLBs were essentially identical to purified hIgGs (Fig. 6 b-c, e, SI Fig. 18a-d, SI Fig. 19 a-b) indicating functional CP6-passivation of SLBs. To further test the efficacy of CP6-based passivation, we compared the effect of lysate on bound mass density for hIgG1 at 1 and 10 nM, and observed no significant effect on lysate-containing sample, indicating that CP6-passivation prevented non-specific protein binding while minimally influencing specific protein-protein interactions (Fig. 6f, SI Fig. 20).

**Figure 6.**
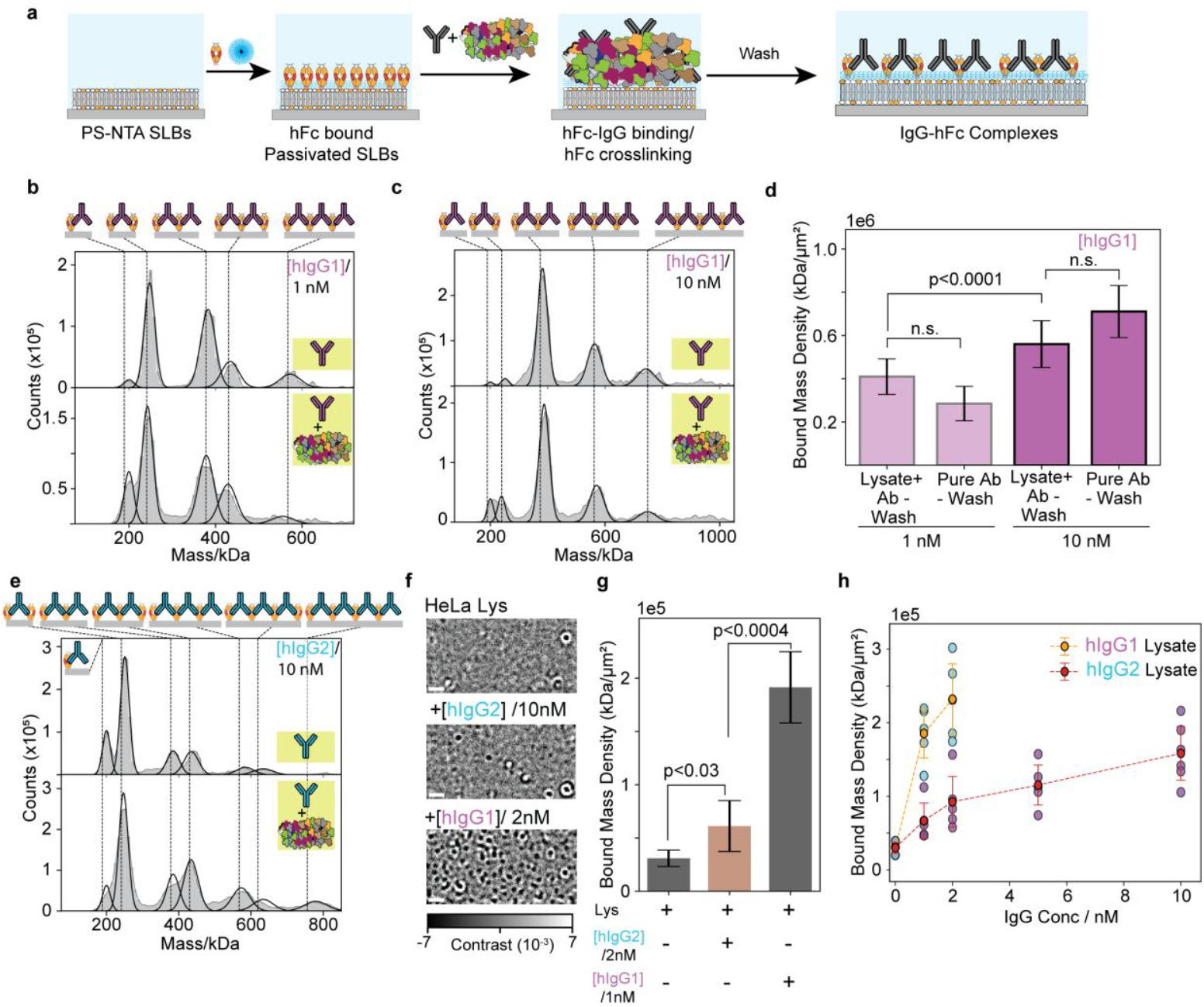
Single-molecule antibody detection and affinity differentiation from mammalian cell lysate. a) Schematic of antibody complex formation in cell lysate. b-c) Comparison of hFc-hIgG1 and hFc-hIgG1 mixed with 0.2 mg/ml HeLa cell lysate on CP6 passivated PS-NTA SLBs containing membrane-bound hFc after 5 minutes of incubation and buffer wash. For each dataset, four 1-minute movies were analysed and pooled. d) Quantification of hIgG1 pulldown at 1 and 10 nM, in the absence (purified) or presence of lysate. We evaluated statistical significance using Mann–Whitney U test. e) hIgG2 binding in the presence and absence of HeLa lysate. For each dataset, 4, 1-minute movies were pooled plotted. f) Representative images of HeLa lysate and HeLa Lysate in the presence of hIgG1/2 showing particle binding in the presence of HeLa lysate, or HeLa lysate containing hIgG1/2. Scale bar: 1 µm. g) Quantification of bound mass in the presence of HeLa lysate, hIgG2+HeLa lysate and hIgG1+HeLa lysate. Both hIgG1 and 2 showed higher binding than HeLa lysate alone. h) Affinity driven single-molecule detection for hIgG1/2 in the presence of HeLa lysate.

To further test the suitability of CP6-passivation for single molecule detection in lysate, we next measured affinity-based detection of hIgG1/2 in the presence of lysate. CP6-passivated bilayers containing membrane-bound hFc were exposed to 0.2 mg/ml HeLa lysate. HeLa lysate alone showed some bound density, likely caused by transient particle binding (Fig. 6f, HeLa Lys; SI movie 9), resulting in a similar visual density to hlgG2 at 10 nM. Addition of hlgG2 at 2 nM or hIgG1 at 1nM, however, showed clear additional binding (Fig. 6g; SI movie 10-11). The difference in bound antibody scaled with the affinity and remained consistent across the concentration range. Accordingly, the trend of total mass with IgG concentration observed without lysate (Fig. 5g-h), was reproduced in the lysate background (Fig. 6h).

## Discussion

We have presented mass photometry-based single-molecule pulldown that enables, mass measurement, abundance and stoichiometry quantification directly from complex mixtures. Our approach expands the scope of MP from purified samples, paving the way for measurement of endogenous protein complexes directly from cell lysates. We combined the intrinsic passivation of lipid surfaces with CP6-based PEG coating to develop a passivated surface that prevented non-specific binding from a variety of cell lysates and serum, indicative of comprehensive passivation compared to previous approaches that were optimized against a single source of non-specific binding^4,9,32,56–59^. Our surfaces can be prepared quickly, and require less standardization of PEG chain length, density, and surface chemistry than PEG coating of bare glass^58–61^. Crucially, CP6-based passivation enables removal of non-specific proteins by simple washing with physiological buffer, essential to preserve low affinity interactions and multi-protein complexes.

The single-molecule mass readout of our assay provides molecular resolution, enabling the identification and quantification of different protein complexes in a single measurement, which is challenging with existing single-molecule pulldown platforms^62^. We demonstrated quantification of protein concentration and oligomeric state straight from *E*.*coli* lysate with as little as 1 µl of clarified lysate, or ∼2×10^5^ cells (see Methods for calculation). Crucially, both the recruitment to the SLB and the presence of cell lysate preserve the oligomeric distribution observed for purified protein in solution. The ability to detect expressed proteins and quantify their oligomeric distribution from lysate removes the need for laborious purification and sample characterization, thereby accelerating the time from expression to experiment. Using hFc-hIgG as an antigen-antibody system, we demonstrated affinity estimation, detection of antibody-mediated antigen crosslinking and stoichiometry. Compared to the bulk readout in BLI and ELISA, resolving all participating species participating allows us to account for avidity effects and accurately determine affinities and antigen crosslinking.

Current limitations are lower sensitivity (∼150 ng/ml) (1 nM) compared to sub ng/ml levels attainable by ELISA, which can be addressed in the future by optimisation of capture complex density and incubation time, opening the door for studies of low abundance proteins^63,64^. Furthermore, SLB-based detection currently exhibits a detection limit on the order of 100 kDa, which can be addressed with capture complexes designed to have a mass >100 kDa. We envision that our platform can become tool for protein detection from biological fluids and cell lysate and become integrated in analytical toolkit of biopharmaceutical companies producing and screening therapeutic antibodies.

## Material and Methods

### Reagents, buffer compositions and instruments

Unless otherwise stated, reagents source and catalogue number were the same as used in earlier study^26^. Assay buffer: 20 mM HEPES (pH 7.45), 150 mM KCl, and was degassed before experiments. Fusion buffer: 50mM Tris (pH 8), 150 mM NaCl+2 mM MgCl2. Ni buffer: Assay buffer + 100 µM NiCl2. Passivation buffer: Assay buffer+ 20 µM CP6. DTT stock: 1M DTT stock was prepared in degassed MILLIQ^TM^, aliquoted and stored in -20°C. Storage buffer: assay buffer + 10% Glycerol. Cell lysis buffer: storage buffer + 1x Protease inhibitor cocktail (PIC) (Roche: 4693132001). 10x PIC solution: 1 PIC tablet was dissolved in 5 ml assay buffer and used as 10x PIC stock. The 10x PIC solution was stored at -20°C until use.

### Protein, chemicals, and reagents for mass photometry

Dyn1-ΔPRD (Dyn1) was expressed and purified as described earlier^1^. Briefly E. coli cells expressing Dyn1 were lysed using fluidizer and lysate was spun at 20000g for 20 minutes at 4°C and the supernatant was collected for Dyn1 purification. Part of the supernatant was aliquoted and flash frozen and was later used as E.coli lysate containing over-expressed Dyn1 (Fig.3) to pulldown Dyn1. To purify full-length protein from the remaining supernatant, nickel affinity chromatography was used, followed by a strep-trap purification. The purity of full-length Dyn1 was tested using SDS-PAGE (Fig. 2a), and protein was flash frozen and stored at -80°C until use.

### Mass photometry experiments

#### Protein preparation

To obtain the solution distribution of Dyn1, Dyn1 stock was dialysed overnight in degassed assay buffer containing 1mM DTT. After dialysis, protein was spun at 20,000 g for 20 minutes at 4°C and the supernatant was collected. The protein concentration was estimated based on the OD_280nm_ (ε_280_= 54,300 M^-1^cm^-1^).

Lyophilised Bovine Serum Albumin (BSA) (Merck, A2153) was used to prepare fresh solution for experiments. Stock was diluted to required concentration and used immediately.

Human IgG1 Fc Protein (AcroBIOSYSTEMS; IG1H5225) was purchased, dissolved in storage buffer at 7.5 µM, aliquoted, flash frozen and stored at -80°C.

hIgG1(Abcam; ab125912), hIgG2(Abcam; ab1927), and mIgG(Abcam; ab125913) antibodies were stored as per the manufacturer’s instructions. For short-term storage (<month), antibodies were stored at 4°C. Antibodies stocks were diluted in assay buffer and used on same day.

#### Mass photometry data Acquisition and movie processing for solution measurement

For mass photometry measurements, glass coverslips (24 × 50 mm, Menzel Gläser) were cleaned using sonication for 5 minutes, first in 50% isopropanol in (Milli-Q® water, 18.2 MΩ·cm), followed sonication in Milli-Q®, followed by drying under constant nitrogen stream. Silicon gaskets (Grace Bio-Labs, GBL103250) were used generate chambers for landing assay experiments.

To obtain solution distribution of Dyn1, Dyn1 stock was diluted to required concentrations, incubated at room temperature for 5 minutes and measured on a mass photometer (OneMP, Refeyn Ltd.).

To measure the mass distribution of pure BSA in solution, freshly prepared BSA stock was diluted in assay buffer to 25nM and used immediately. For BSA + E. coli lysate measurements in solution, BSA stock was diluted to 25 nM in 0.2mg/ML E.coli lysate (see below for lysate(s) concentration estimation) and measured.

Movies were acquired with Acquire MP at 360 Hz, and movie processing and particle detection was carried out using DiscoverMP version 2023 R1.2 (Refeyn). All analysis parameters were left at the default values (5 frames averaging, threshold 1: 1.5, threshold 2: 0.2, motion correction and reflectivity correction enabled). Contrast to mass conversion was applied as mentioned earlier^17,26,29,65^. Dyn1 was used as contrast to mass calibration and has been shown to match other mass calibrants^26^.

For oligomer distribution for purified BSA, and BSA + E. coli lysate, mass histograms (5.6 kDa bin width) were generated with DiscoverMP. For Dyn1 solution distribution, the list of events was exported as CSV file for further analysis in custom Python scripts. Mass histograms were generated (Numpy.histogram, 10.5 kDa bin width, mass range 0-670 kDa). The distribution was fitted (scipy.optimize) with a mixture of Gaussians.

#### hFc and hIgGs affinity measurement

To perform affinity measurements for hFc and hIgG, we were limited by antibody binding to glass and found that at 5 nM, hIgGs provided optimal, mass resolved histograms (Fig. S13). For hFc-hIgG affinity measurements, 5 nM hIgG was incubated with increasing hFc at room temperature. For hIgG1, 10 nM hFc was sufficient to form hIgG1-hFc complexes almost immediately (Fig. S13). However, for hIgG2, we had to increase hFc concentration to up to 45 nM and incubate for 20-30 minutes to see significant formation of hIgG2-hFc complexes (Fig. S13). Data was analysed with Discover MP as described before.

### Dynamic Mass photometry measurement and movie processing

#### Liposome preparation

##### PS-NTA liposomes

Liposome preparation was performed as described earlier^26^ with minor modifications: For PS-NTA liposomes, DOPC, DOPS, and DGS-NTA lipids were mixed at 89:10:1 ratio. Required volume of chloroform stock of DOPC, DOPS and DGS-NTA were added to pre-cleaned glass vial at the required ratio to obtained 500 µM total lipid concentration. Excess chloroform was removed via rotary evaporation under constant nitrogen stream. The resulting lipid-film was hydrated in 500 µl assay buffer for 30 minutes in 50°C water bath with intermittent mixing. Subsequently, liposomes were transferred into 1.5 ml Eppendorf tubes and sonicated (Vibra-Cell VC505; Sonics) using 3 mm probe at 25% amplitude with 1 sec on-3 sec off sonication cycle. Sonication was done in a beaker containing ice-water mix to ensure efficient transfer of the heat generated during sonication. Sonicated liposomes were spun at 20,000g for 30 minutes at 4 degrees and supernatant was collected in a clean Eppendorf tube, stored at 4°C and used within a week.

##### PEG550-PS-NTA liposomes

DOPC, DOPE-PEG550, DOPS and DGS-NTA were mixed at 79:10:10:1 mol% to obtain 500 µM total lipid concentration. DOPE-PEG550 stock was prepared in chloroform, required lipid stock for each lipid was aliquoted in a pre-cleaned glass vial and dried via rotary evaporation to generate thin lipid films. Liposome hydration and sonication procedure was the same as for PS-NTA liposomes.

#### Supported bilayer (SLB) deposition

For bilayer deposition, glass coverslips (24×50 mm, Menzel Gläser, VWR 630-2603) were cleaned by sonication in an ultrasonic bath in 50% isopropanol for 5 min, followed by sonication in Milli-Q for 5 min, and dried using a nitrogen stream. For SLB preparation, coverslips were activated via plasma treatment (Zepto2, Diener Electronics GmbH) in the presence of oxygen at 40% power for 4 minutes and used immediately. For bilayer deposition, 30 µl of fusion buffer was added in a silicon gasket (Grace Bio-Labs, GBL103280-10EA) and 20 µl of 500 µM sonicated liposomes were mixed in. Bilayer formation occurred by liposome fusion to the glass surface and was confirmed with the mass photometer by observing the formation of a homogenous film. Bilayers were incubated at RT for 30 minutes and excess liposomes were washed with 10 ml (1 mlx 10) assay buffer. 20 µl volume of 3 mm gasket was supplemented with 40 µl buffer to generate 60 µl reaction volume. To charge NTA lipids with Ni, 40 µl assay buffer was replaced with 40 µl Ni buffer for 5 minutes and washed via serial dilution, 40µl x 5 times (200 µl), with assay buffer and stored in humidity chambers till use.

#### Bilayer passivation, and single-molecule Dyn1 pulldown

##### Bilayer mediated passivation against BSA

To test the passivation against non-specific protein binding, PS-NTA SLBs were treated with increasing BSA concentrations. To achieve these concentrations, BSA stock was diluted to 1.5x the desired concentration and 40 µl of 60 μl of gasket volume was replaced with 40 µl of 1.5x stock of desired BSA concentration and incubated for 5 minutes. To test the effect of protein incubation time on non-specific binding, two strategies were tested; 1) short incubation, where after incubation with desired BSA concentration for 5 minutes, bilayers were washed using with 300µl assay buffer (Fig. 1g), and long incubation, where bilayers were exposed to increasing BSA concentration for 5 minutes, multiple field captured, and next BSA concentration was added (SI Fig. 3). After measuring highest BSA concentration (100µM BSA), bilayers were washed with 300µl assay buffer. Both strategies showed similar results indicating that a bilayer’s ability to prevent non-specific binding is not affected by the exposure time.

#### Washing unbound proteins via serial dilution

The gaskets used for SLB measurements require between 20 μL and 60 μL sample volume. After an incubation of 60 μl sample, we wash by removing 40 µl of sample and replace it with required volume and repeated the dilution multiple times. In the paper, this washing process is referred as “*V*×n” where *V* represents the volume of buffer and *n* indicates the number of times the dilution was repeated. For example, 300 µl (100 µl×3) wash indicates that the gasket volume (∼20 µl) was diluted with 100 µl buffer, diluting content 6 times, and 3 repeats of washing effectively dilute the content 6^3^,(216 times), finally, 40 µl buffer is added to make 60 µl reaction volume before imaging, thus further increasing dilution by 3 fold, effectively diluting solution concentration of proteins by 648 times. Hence, for wash used in the experiments, the effective dilution can be obtained using this formula. (((*V*+20μl)/20 μl)^*n*^)*3, where the x is the volume used for serial dilution and *n* is number of times the serial dilution was repeated.

##### Single-molecule Dyn1 pulldown from BSA-Dyn1 mixtures

For Dyn1 pulldown from Dyn1+ BSA mixture, Dyn1 at increasing concentration was premixed with fixed BSA concentration and incubated with PS-NTA bilayers. For single-molecule pulldowns, minimising Dyn1 concentration loss, especially at low Dyn1 concentration (<100 nM) is critically important. We used high BSA concentration to minimise Dyn1 loss to the walls of the tube. For pulldown experiments, we incubated 0.1, 1 and 2.5 nM Dyn1 mixed with 100 µM BSA, and to achieve these final Dyn1 and BSA concentrations, we prepared 1.5× the dyn1+BSA mixture concentrations and added 40 µl of the mixture to the gasket containing 20 µl assay buffer, mixed and incubated. To prepare target concentrations, we prepared 150 nM Dyn1 stock in 150 µM BSA and serially diluted in 150 µM BSA to achieve 0.15 nM Dyn1+150 µM BSA, 1.5 nM Dyn1+150 µM BSA, and 3.75nM Dyn1 + 150 µM BSA. Subsequently, SLBs were incubated with higher Dyn1 concentration and the procedure repeated until all concentration were tested. Three repeats with separate bilayers were performed.

#### Dyn1 pulldown from E. coli lysate

For Dyn1 pulldown from E. coli lysate, flash-frozen E. coli supernatant was thawed on ice and added to PS-NTA bilayers. E. coli lysate was diluted to 50% (lysate stock) and used as a stock for Dyn1 pulldown. First, we added 1 µl lysate stock to 60 µl assay volume of PS-NTA bilayers (0.0083%), incubated and measured without washing excess lysate. Subsequently, we added increasing lysate concentration and measured the emergence of oligomers (Fig. S4). After testing highest lysate concentration, bilayers were washed and the Dyn1 oligomeric distribution was measured (Fig. S5). To test the efficacy of the SLB passivation against E.coli lysate proteins, 10% E.coli lysate on separate SLB was added, incubated for 5 minutes, washed and imaged (Fig. 9).

We initially had ∼1 x 10^9^ cells per mL in the culture (based on OD600 of 1), so 1 x 10^12^ total cells in our 1 L culture^66^. Cells grown in 1L lysate were pelleted, lysed into 65 ml final volume. We use ∼100 uL measurement volume and dilute 1.2 x 10^4^ fold (0.0083%). So, we effectively use: 1×10^12^ x0.1 / 65 / (1.2*10^4^) = 1 x 10^5^ cells (which corresponds to < 0.1 µl of our total cell culture volume).

#### Cholesterol-PEG600 (CP6) solution preparation and bilayer passivation

Cholesterol-PEG600 (Avanti polar lipids; 880001O) oil solution was dissolved in anhydrous methanol at 50 mM stock and stored at -20 °C in a clear, glass vials with PTFE liner cap, secured with parafilm. For bilayer passivation, 1 mM CP6 stock was prepared in assay buffer and stored at 4 ^°^C till use. This stock was used to obtain the mass histogram for CP6 at 20 µM (Fig. 4b). For bilayer passivation, two strategies were tested: Direct dilution: 1mM stock was diluted into the gasket at final concentration of 20 µM; serial dilution: we made 20 µM stock in assay buffer and washed PS-NTA SLBs by replacing 40 µl buffer with 20 µM CP6 stock and repeated the dilution 5 times to achieve 20 µM CP6 concentration in the reaction chamber. In both cases, CP6 was incubated for 5 minutes and then washed with assay buffer to remove excess CP6 and provided similar passivation levels. Serial dilution was used for pull down experiments.

To test whether CP6 passivation affects single molecule pulldown, on CP6 fortified PS-NTA bilayers, we added 0.016% E.coli supernatant and incubated for 5 minutes and collected oligomer diffusion without washing the lysate (Fig. S12).

#### Mammalian cell lysate preparation, and CP6 passivation efficacy against lysates

##### Cell lysate preparation

Mammalian cells (SHSy5y, HEK, HeLa) were resuspended to 1 mil/ml and 10 ml of cell suspension was spun at 500g at 4°C for 10 minutes to minimize mechanical disruption. Supernatant was discarded and cell pellet was resuspended in 10 ml PBS and spun down at 500g at 4°C for 10 minutes. The process was repeated 2 more times and cell pellet after final wash was resuspended in 1 ml ice-cold cell lysis buffer and lysed over ice using mechanical disruption using Dounce homogeniser. 25 strokes were used to ensure complete lysis. The lysate was spun at 20,000g for 30 minutes at 4 °C to remove insoluble debris. Supernatant was aliquoted, flash frozen in liquid nitrogen and stored in -80 °C until use. BCA assay (ThermoFisher; A55864) was used to estimate the total protein concentration in cell lysates, and concentration estimated around 2mg/ml for all lysates.

##### PEG550 mediated passivation against cell lysate

To evaluate lysate passivation by PEG550 lipids, we used PEG550-PS-NTA liposomes to make pegylated SLBs. For SLB preparation, 20 µl liposomes (500 µM stock) was mixed in gasket containing 30ul of fusion buffer over a plasma cleaned coverslip and incubated for 30 minutes. After washing excess liposomes bilayers were used for testing passivation capacity. PEG-SLB was incubated with 2e-1 mg/ml SHSy5y lysate, incubated for 5 minutes at RT, and washed with assay buffer. Multiple fields of view were imaged.

##### CP6 mediated passivation against cell lysates

To test the efficacy of the CP6 passivation against different cell lysates, we fortified PS-NTA bilayers with CP6 as described earlier and then exposed to increasing cell lysate concentrations. For each cell lysate, we exposed SLBs to 2e-3, 4e-3, 2e-2 and 2e-1 mg/ml lysate and collected movies of different fields. After exposure to highest lysate concentration, SLBs were washed with assay buffer, 100µl ×3, and imaged. For bilayer exposed to Foetal bovine serum (ThermoFisher; A5670401), we needed to carry out excess washes to remove bound particles 1.5 ml (100 µl x 15).

#### hFc-migG cross-reactivity testing

To test cross-reactivity between hFc and non-specific mIgG, we added 2 nM hFc on CP6-fortified PS-NTA bilayers. Without washing excess hFc, increasing concentration of mIgG up to 100 nM was added to the bilayer. At 50 times higher concentration than the hFc, we did not observe any binding (Fig 5c). Under these conditions, increasing presence of hIgG2 led to complex formation (Fig. S14), showing the specificity of hFc-hIgG2 interactions.

#### hFc-hIgG interaction stoichiometry and affinity determination

To estimate hFc-hIgG affinity, we titrated hIgG1/2 concentration over a bilayer containing fixed hFc concentration. To prepare hFc bilayers, CP6 fortified PS-NTA bilayers were incubated with 2 nM hFc for 5 minutes and excess hFc was washed via serial dilution (40µl ×3) giving effective hFc solution concentration to 74 pM. Subsequently, increasing concentration of higG1or hIgG2 was added to these SLBs. For each concentration, 1-minute movie from 5 separate fields were acquired. Movies were acquired at 270 Hz.

#### hIgG detection from lysates

To detect antibodies from lysate, we hFc treated bilayers (as described above) with HeLa lysate (0.2mg/ml) and hIgG stock was diluted to reach required concentration and bilayers were incubated for 5 minutes at RT, washed (Fig. 6 a) and 1-minute movies of different fields were acquired.

#### Data analysis for dynamic mass photometry

Dynamic MP measurements were carried out on oneMP mass photometer (Refeyn Ltd). Movies were acquired at maximum frame rate of 360 Hz on a 9.9 x 9.9 µm^2^ FOV at an effective pixel size of 77.34 nm after 4×4 pixel binning. All movies were collected for 1 minute.

#### Image processing

was carried out as described previously^26,29^ with a few modifications. For image processing, movies were processed as described earlier^26^ with the exception of no frame averaging, thus the final frame rate remained 360 Hz. To remove static glass background (roughness), images were processed through sliding median ratiometric background removal^26,29^. For this, a predefined frames window centred around the frame of interest was used to calculate the median background (100 frames either side of the raw frame of the interest, 201 frame or 609 ms) and used to remove the static glass background by dividing the raw frame by the background median image^1,2^ and subtracting 1.0, revealing particles diffusing on the bilayer.

#### Particle detection

For particle detection, Laplacian of Gaussian kernel with a sigma of 1.5 pixel was convoluted with each mobile frame, followed by application of a manual threshold 0.0085 to generate a binary map of particles contrast exceeding this threshold.

#### Particle fitting

For each detection, a 15×15 px thumbnail centred around the detected position is extracted. The thumbnails are fitted with an analytical PSF model^26^ using a gradient descent optimiser to minimise the mean squared error. The fitting procedure is accelerated by pre-calculation of the PSF on a sub-nm grid and using a look-up instead of calculating the PSF for every fit iteration. The fit returns a sub-pixel event position, event contrast, and fit residuals.

#### Trajectory linking and diffusion estimation

To determine the diffusion coefficient, we linked 2D particle localizations across successive frames using their x,y positions to generate trajectories, which were analysed to extract diffusion coefficients. Particle linking was performed with Trackpy’s link_df function, as previously described^26,29,^ The linking function has 4 relevant input parameters: a) The maximum displacement (*d*max) was calculated using the previously described equaltion^29^; assuming a maximum diffusion coefficient 1.5 µm^2^/sec and time between consecutive frames of 0.003 s (340 Hz), yielding a maximum expected displacement of 0.346 µm between consecutive frames, corresponding to 5 pixels. The linking memory was set to 2 frames to enable linking across missing detections. Prior to linking particles with a contrast below 0.003 (≈ 100 kDa) were removed.

Only trajectories lasting at least 30 frames; 90 msec (for data in figure 2-3) and 40 frames; 120 msec (for data in 5-6) were selected for further analysis. Trackpy-based linking does not account for particle contrast and particle movement can lead to merging trajectories or trajectories that cross path with each other, which can lead to aberrant contrast. To remove these noisy and poorly linked trajectories, standard deviation of the contrast was used as described earlier^1^. Next, we constructed a contrast histogram for each trajectory and applied a Gaussian fit to extract the mean and standard deviation of the contrast of each trajectory^26^. Subsequently, mean + 2 s.d. was used to ascribe trajectories to a particular mass to an oligomer (Dyn1) or a protein complex (hFc-hIgG). Finally, diffusion coefficients for each trajectory were calculated as described earlier^1^ and we observed that more than 95% particles displayed single mobility component.

#### Dyn1 concentration estimation from E. coli lysate

##### a) Bound mass density in dependence of Dyn1 concentration

To estimate the concentration of over-expressed Dyn1 in E. coli lysate, Dyn1 oligomer binding to PS-NTA bilayers containing a limiting number of binding sites provided by DGS-NTA lipid (NTA lipid) in the bilayer was measured. Because the oligomeric state of Dyn1 sensitively depends on its concentration, increasing Dyn1 concentration simultaneously leads to more bound complexes and a shift towards larger order Dyn1 oligomers, at the expense of smaller ones (Fig. 2e). To account for protein binding and change in oligomeric abundance, we devised a bound mass density (kDa/µm^2^) (BMD) to reflect total mass bound to the bilayer regardless of oligomeric state: From the trajectory linked data, trajectory-length weighted histograms over the 0 to 850 kDa mass range were generated and the peaks of all oligomeric species fitted with a mixture of Gaussians. To estimate the BMD, the integral of each Gaussian component was multiplied by its expected mass and summed. The resulting sum was normalised to the measurement duration and detection field of view. As calibration, the BMD was measured for a titration of Dyn1 in presence of 100 µM BSA. To confirm the direct scaling of BDM with concentration a linear fit without offset was performed (Fig. 3c).

##### b) Dyn1 concentration estimation in complex mixtures

The BMD to concentration relation was used to extract Dyn1 solution concentration from separate Dyn1+BSA experiments (Fig. 3d). To estimate Byn1 concentration in E. coli lysate, we extracted BMD for each lysate dilution and used the Dyn1-BMD to concentration relation to extract Dyn1 concentration at each lysate dilution (Fig. 3e).

#### Bound mass density estimation for hFc-antibody binding

For hFc-hIgG interaction mediated hIgG recruitment to the CP6 fortified NTA bilayers, the density of multiple species was poor fit for gaussian fitting based area calculation (Fig. 5f (5nM), 5g (50nM); Fig. S14). To ensure we can accurately estimate the area for all species observed in our data, we adopted an alternative approach to use nonparametric means of area calculation. We defined a mass range (0-1500 kDa) and bin width (10.5 kDa), weighted all mean trajectory mass with its trajectory length, and generated a frequency distribution. We next used interp1d from scipy.interpolate to generate cubic interpolation of bin-centres and respective frequency distribution and used quad function from scipy.integrate to calculate the mass weighted integrated area. The resultant area was then normalised to the movie length and the area of the acquisition to generate bound mass density (kDa/µm^2^).

#### hIgGs affinity to hFc extraction

To measure the hIgG1/2 binding affinity to hFc, we performed a hIgG1/2 titration on bilayers with a fixed hFc concentration. The extracted bound mass density (kDa/µm2) reflected the increase in hIgG bound to the hFc containing bilayers, and resembled the single site binding curve (Langmuir isotherm). To extract affinity, we fitted all data points to both single site binding and hill equation (eq.1). We observed data fit better to hill equation (R2 ≥ 0.9) indicating that on bilayers, both hIgGs showed cooperativity to bind 2-hFc (Fig. 5g-h), and hIgG1 showed relatively higher cooperativity than hIgG2 (Fig. 5 g-h), indicating a higher potential to crosslink hFc on bilayers. Our data also showed that hIgG1 showed 8-10 times higher affinity than hIgG2 (Fig. 5 g-h).

#### hIgG detection in lysate

To test the efficacy of hFc-hIgGs interaction to on CP6 passivated bilayers is affected by the presence of mammalian cell lysate, we estimated bound mass density before and after lysate wash. To test this we incubated 1, and 10 nM hIgG1 on CP6 fortified hFc bilayers for 10 minutes, captured distribution across multiple fields and washed with 300ul buffer (100µl ×3) diluting the hIgG1 concentration by 216 (6^3^) times leading to effective concentration to 4.6 and 46 pM, respectively. We observed no change in density due to washing (Fig. S21). Next, we tested the effect of the presence of lysate and compared bound mass density for 1 and 10 nM hIgG1 with and without HeLa cell lysates (0.2mg/ml). Incubation, followed by wash, did not significantly affected the bound density for hIgG1 at either concentration (Fig. 6f).

To further quantitatively compare whether we can detect specific binding in the presence of lysate, we prepared two, CP6-fortified bilayers, treated with 2nM hFc for 5 minutes, washed excess, unbound hFc, bilayers were treated with 0.2mg/ml HeLa lysates. Subsequently, one bilayer was exposed to increasing hIgG1 concentration, and another with hIgG2 concentration. For each concentration, we collected at least 5, 1-minutes movies after hIgG1/2 addition, before the bilayers were treated with the next hIgG concentration. After collecting highest concentration for each hIgG, we washed bilayers with 300µl buffer and collected 5 more movies of different field of views. For each concentration, then we plotted bound mass density as a function of hIgG1/2 concentration.

#### Effect of non-specific binding on Dynamic MP image noise calculations

The indiscriminate, non-specific binding of BSA (Fig. 2g, 3c), or binding of large number of proteins with unknown mass in case of cell lysates (Fig. 4), many below the detection limit of dynamic MP, 120 kDa, makes it difficult to accurately quantify the mass of the bilayer bound protein. To understand how indiscriminate protein binding to the bilayer affects dynamic MP signal and impeded detection of a specifically bound protein, we used standard deviation of the background subtracted dynamic MP image. Image noise divided by the contrast to mass relation provides image noise in kDa, setting the lowest detection limit below which any specifically bound particle has a lower signal than the image noise fluctuations. Image noise/kDa as a function of BSA concentration (Fig. 1g) or cell-lysate concentration led reached a substantial level. However, in both cases, washing with buffer containing physiological salt concentration and pH decreased image noise to the level before exposure, indicating the passivation capacity of bilayers (Fig. 1g) and CP6 mediated bilayer passivation against wide range of cell lysates (Fig. 4e).

#### Trajectory length plot

To plot trajectory length of all Dyn1 oligomers, all particles within mass range (0, 850) kDa were collected and their trajectory length was extracted and converted to seconds (0.0037 sec/frame). The trajectories of the tetramer species were selected based on their mean mass, defined as the expected tetramer molecular weight (360 kDa) corrected for a 5% contrast reduction on bilayers (342 kDa), with an acceptance window of ± 2 SD. Only trajectories meeting this mass criterion were included in the tetramer trajectory-length analysis.

To plot trajectory length for hFc-hIgG1/2 data, all masses within mass range, 0, 1500 kDa were selected.

## Supporting information

SI_figures_compiled

## Acknowledgements

We thank Dr. Tarick J El. Baba and Dr. Di Wu from Professor Dame Carol Robinson’s lab for providing ShSy5y, HEK and HeLa cells. This work was funded by the European Research Council (ERC) Consolidator Grant PHOTOMASS 819593 (P.K., M.S.K., J.C.T.), the Engineering and Physical Research Council (EPSRC) Leadership Fellowship EP/T03419X/1 (P.K., JC.T., E.D.B.F.), the Medical Research Council (MRC) CDT in Biomedical Imaging EP/L016052/1 (E.D.B.F.), St Hugh’s College (E.D.B.F.), the Clarendon Fund (E.D.B. F.), the Biotechnology and Biological Sciences Research Council BB/W00349X/1, (P.K), the Wellcome Trust, Grant Number: 218514/Z/19/Z (R.W), Schmidt Sciences, LLC (J.C.T.).

For the purpose of Open Access, the author has applied a CC BY public copyright licence to any Author Accepted Manuscript (AAM) version arising from this submission.

## Author contributions

P.K., J.L.P.B., M.S.K and E.D.B.F. conceptualised SLB mediated single molecule pulldown with mass photometry. E.D.B.F. carried out single molecule pulldown using SLB. M.S.K and E.D.B.F. carried out Dyn1 pulldown from E.coli lysate. M.S.K. conceptualised CP6 based bilayer passivation. P.K., M.S.K, R.W. and J.C.T. conceptualised antigen-antibody detection from lysates, and M.S.K. and J.C.T designed the antigen-antigen methodology used. M.S.K and R.W. performed comprehensive characterisation of CP6-based passivation against lysates and performed antibody pulldown experiments. J.C.T modified analysis pipeline published earlier^26^. M.S.K. analysed the data. P.K., M.S.K, R.W. and J.C.T. prepared figures and wrote the manuscript.

## Competing interests

P.K. is an academic founder, shareholder, and non-executive director to Refeyn Ltd. All other authors declare that they have no competing interests. J.L.P.B is academic founder, shareholder and consultant to Refeyn Ltd.

