## Supplementary material for "Mass-specific single molecule pull-down from complex mixtures with bilayer-assisted mass photometry": SI_figures_compiled

**Supplementary Figures**

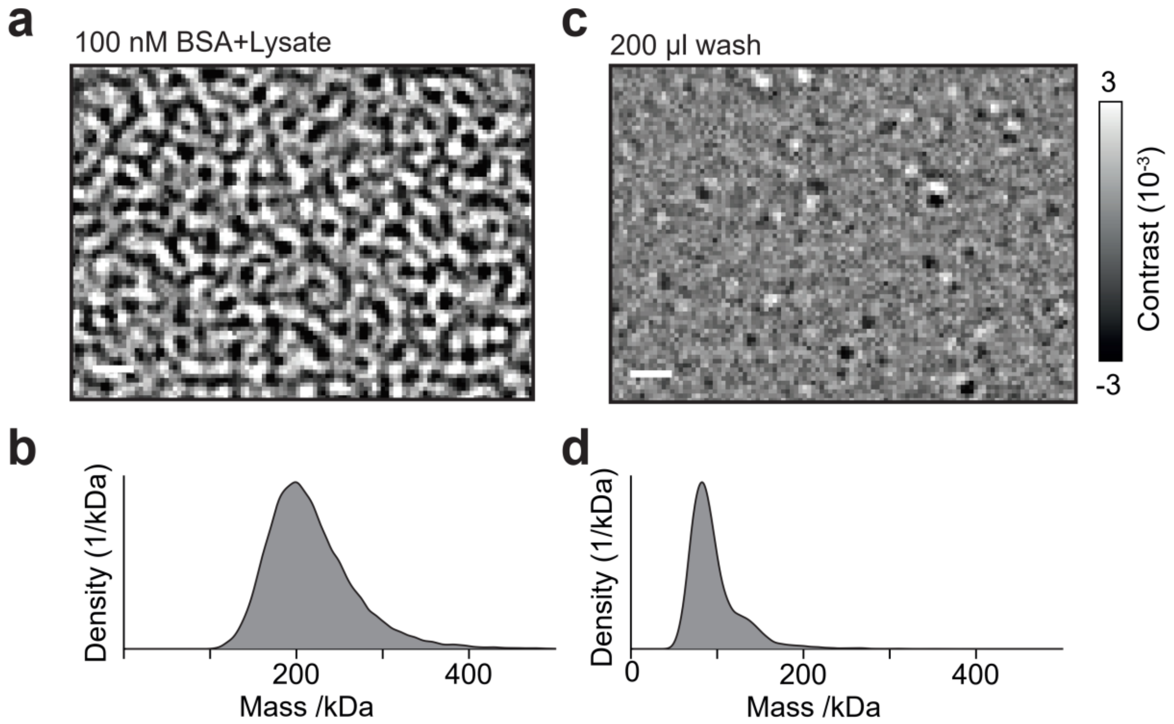


**Supplementary Figure 1**. Effect of lysate on MP mass resolution and ineffficacy of washing unbound protein to obtain resolved mass histogram. a-b) Background subtracted image of the 100nM BSA+E.coli lysate (0.2mg/ml) added to glass coverslips (a) and resulting unresolved mass histogram (b). c-d) Representative background substracted image of glass coverslip after binding BSA and lysate proteins (c), and resulting histogram of bound proteins (d). Due to lack of specific binding sites on glass, the identity of the proteins resulting into the histogram remains unknown.

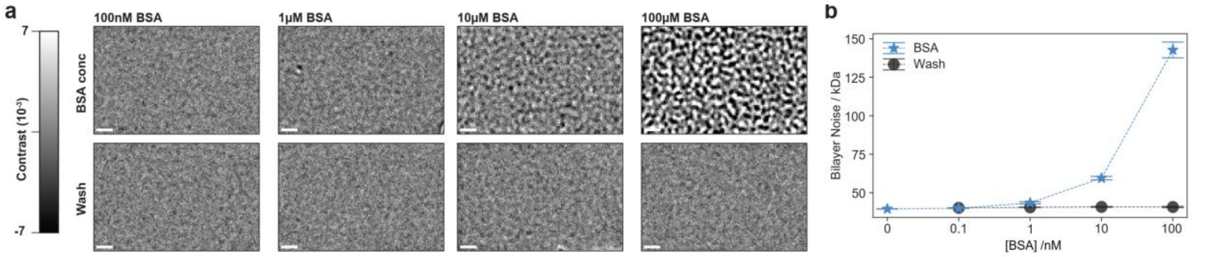


**Supplementary Figure 2**. Bilayer efficacy to prevent non-specific binding. a) Representative median subtracted images of PS-NTA SLBs incubated with increasing BSA concentration (BSA conc) and subsequent wash with reaction buffer (wash). b) Bilayer noise as a function of increasing BSA concentration and subsequent wash. For each condition, 3, 1-inute movies were used to calculate mean and standard deviation of the noise. For 100uM BSA wash, 6, 1 minute movies were used. Noise quantification suggests that SLBs significantly reduce BSA binding to bilayers and at 10μM BSA, marginal shift in BSA binding appears which is reflected both in background subtracted image and bilayer noise increase. Wash data indicates that BSA non-specific binding on bilayers can be removed by washing with buffer containing physiological pH and salt concentrations. Scale bar = 1μm

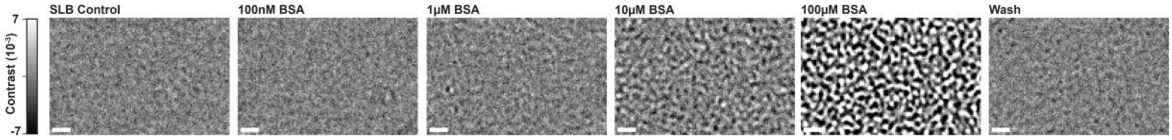


**Supplementary Figure 3**. Effect of BSA addition and incubation on bilayer noise. PS-NTA SLB was incubated with increasing BSA concentration, incubated for 5 minutes at RT and imaged. After 100μM BSA incubation, SLB was washed with reaction buffer. Loss of all non-specifically bound protein suggested prolonged BSA exposure does not decrease passivation by bilayers. Scale bar = 1μm.

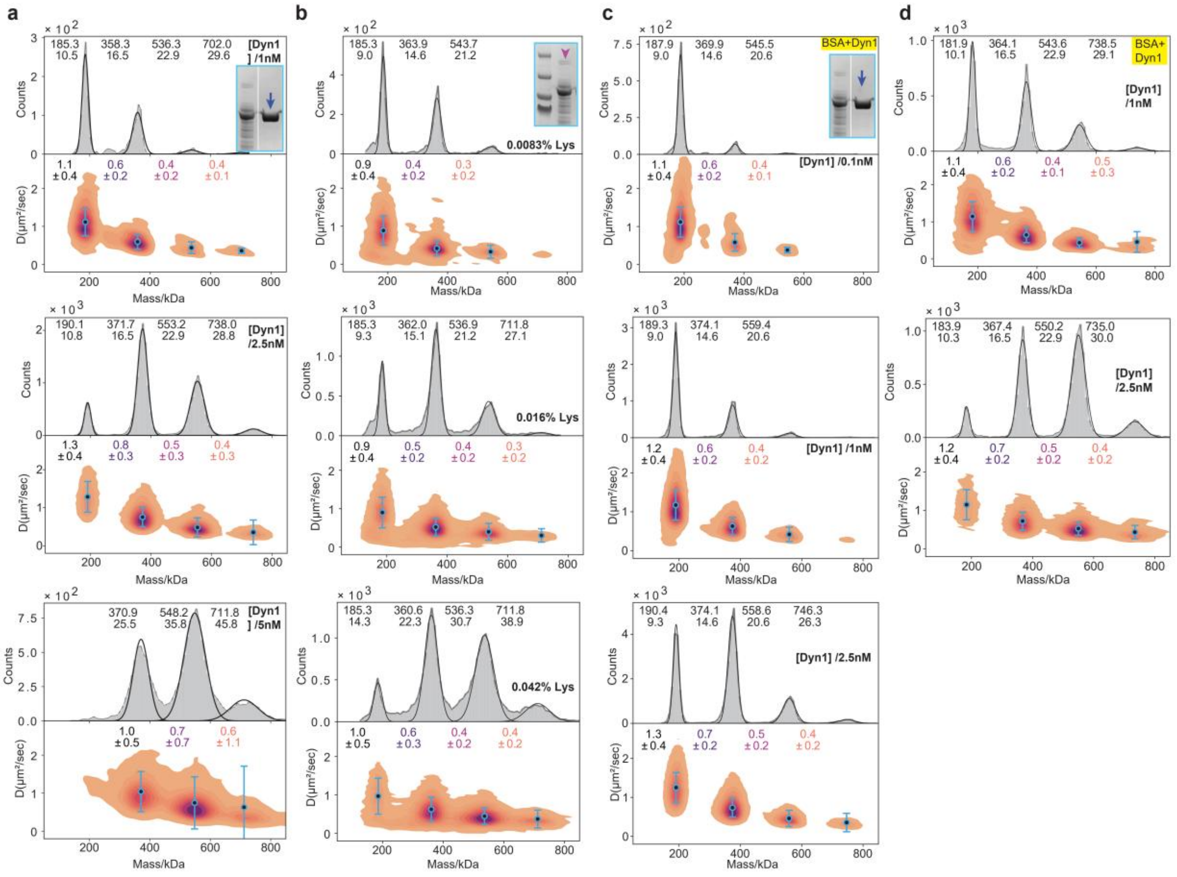

**Supplementary Figure 4**. Mass and diffusion distribution for Dyn1 oligomers recruited on PS-NTA SLBs. a) Mass and diffusion distribution of Purified Dyn1 oligomers as a function of concentration. For 1 and 2.5 nM, data was pooled from 5, 1minute movies. For 5 nM, data from 4, 1-minute movies were pooled. SDS PAGE inset shows the purified protein used in the experiment. At 5 nM Dyn1 solution concentration, binding goes up significantly, leading to the loss of resolution resulting into wider peaks and loss of dimer. Under these high-density conditions, data deviates from linearity and becomes qualitative (see SI Fig. 7 to see the effect of protein density on particle movement on bilayers). b) Mass and diffusion distribution of Dyn1 oligomers obtained from E.coli lysate over-expressing Dyn1 (SDS PAGE inset). Increasing lysate concentration over PS-NTA bilayers leads to similar mass and diffusion coefficient distribution as obtained for purified Dyn1 (a). For each lysate condition, data from at least 6, 1-minute movies was pooled. Similar to 5nM purified Dyn1, addition of highest lysate concentration (0.042%) led to peak broadening and loss of peak resolution. c-d) Dyn1 oligomer distribution after varying Dyn1 concentration was incubated on PS-NTA SLBs for 5 minutes and unbound protein was washed with reaction buffer. Oligomer in c were used to generate linear relation between Dyn1 concentration and resulting Dyn1 oligomers bound to the bilayer. The linear relation obtained was used to estimate the solution concentration from dyn1 in D. See SI Fig. 5 to correlate Dyn1 peak mass and diffusion to oligomer identity.

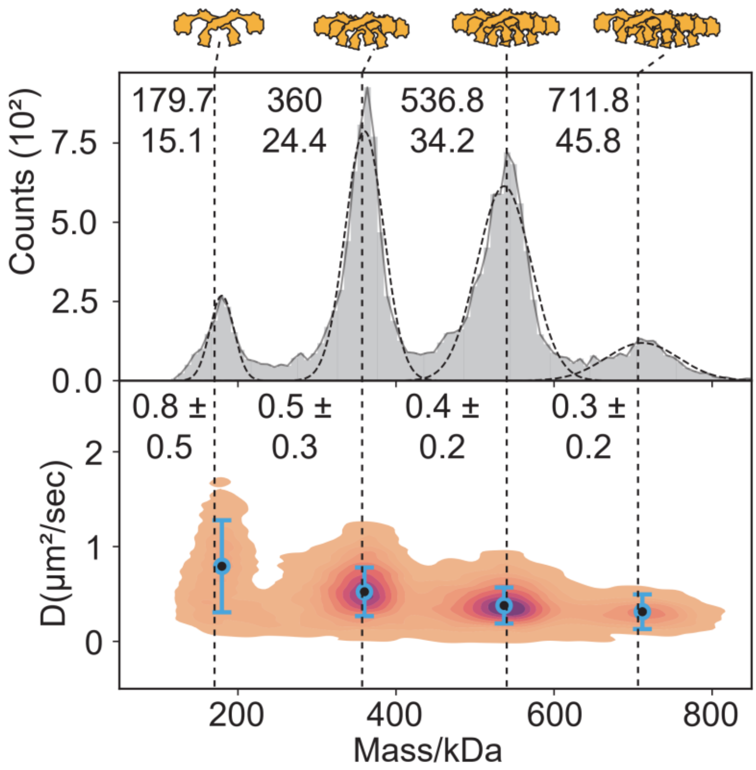


**Supplementary Figure 5**. Mass and diffusion distribution for Dyn1 oligomers on PS-NTA SLBs after wash. After measuring Dyn1 oligomeric distribution at 0.042% lysate concentration, SLBs were washed with 500ul (100 ul x5) assay buffer leading to the final lysate concentration in solution to be infinitely small ( 5.4e-6%; see methods for details). Data from 7, 1 minute movies (n=7) was pooled and used to plot the masss and diffusion histogram.

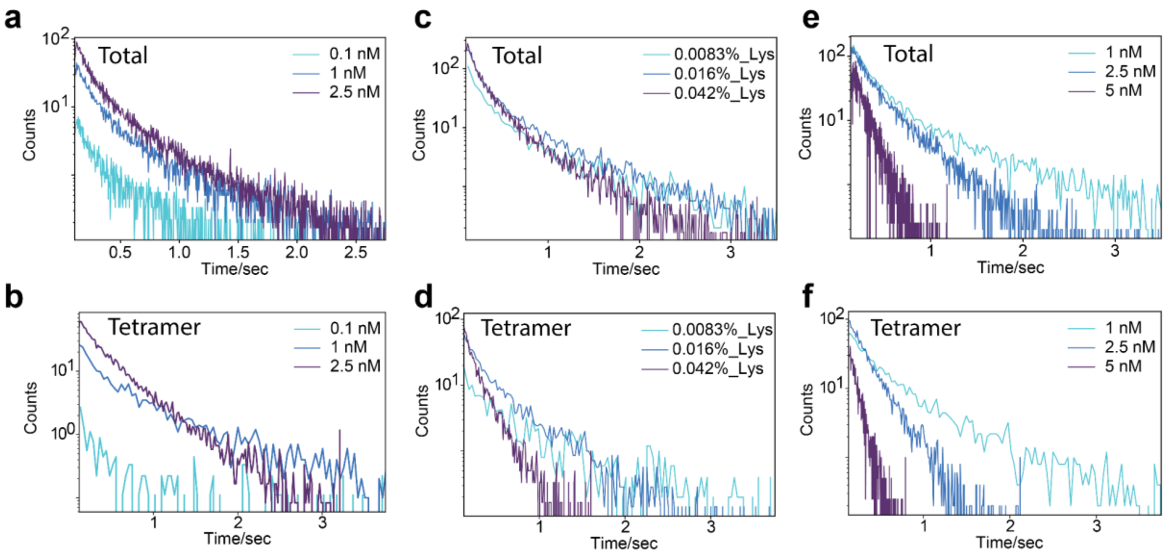


**Supplementary figure 6**. Trajectory length distribution as a function of particle density on bilayers. a-b) Trajectory length for all Dyn1 oligomers (a) and tetramer (b) for Dyn1 titration in 100 μM BSA (SI Fig. 4) was combined and plotted. c-d) Trajectory lengh for all Dyn1 oligomers (c) and tetramer (d) pulled down from E.coli lysate dilutions. e-f) Trajectory length for Dyn1 oligomers (e) (purified) and tetramer (f) from Dy1 (purified) titration on bilayers were plotted. Note the extreme shift in trajectory length change for all oligomer and tetramer in case of high particle density on bilayer (e-f). Short trajectory length for 0.1 nM Dyn1 in case of a-b is because dimer, fastest oligomer is the predominant oligomeric species at 0.1 nM Dyn1 (SI Fig. 4c).

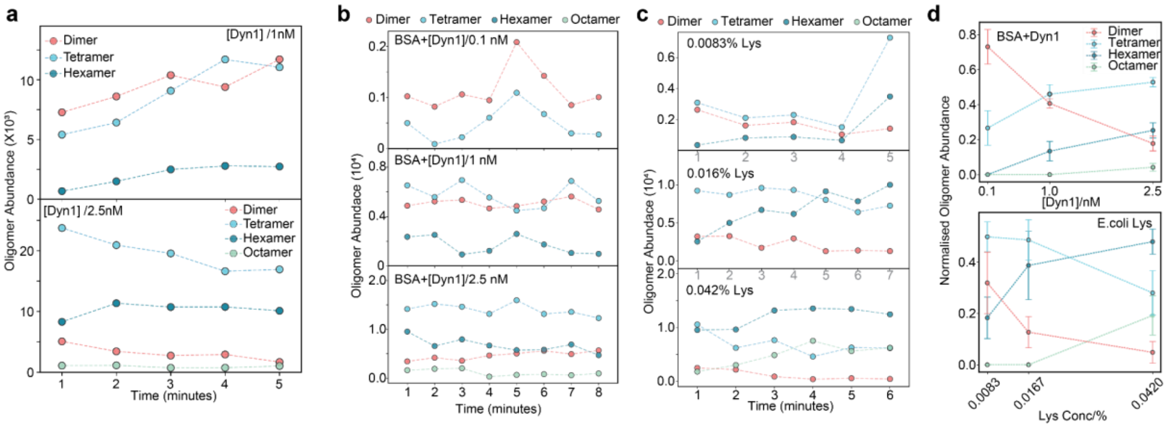

**Supplementary figure 7**. Oligomer abundance for data show in SI Fig.4. a) Dyn1 oligomer abundance for Purified Dyn1 at 1 and 2.5 nM. Data shows change in oligomers over time in different fields of same bilayer. Data also shows the concentration dependent emergence of Dyn1 octamer at 2.5 nM, indicating that Dyn1 oligomeric abundance on SLBs can be influenced by Dyn1 solution concentration. b) Dyn1 oligomer abundance upon incubation of Dyn1+BSA (100 μM) on PS-NTA bilayers. Like purified Dyn1, increase in Dyn1 concentration in presence of BSA leads to emergence of larger Dyn1 oligomers on SLBs. c) Change in Dyn1 oligomeric as a function of increasing E.coli lysate containing Dyn1. Dyn1 oligomeric abundance in lysate mimics similar behaviour to Purified Dyn1 alone or Dyn1 mixed with 100μM BSA. d) Evolution of Dyn1 oligomers as a function of increasing concentration. Comparison of Dyn1 oligomers in the presence of BSA and from E.coli lysate shows similar trend. In both cases, Dyn1 dimer decreases with increasing Dyn1 or lysate concentration. Similarly, increase in the abundance of large oligomers show similar trends, indicating that bilayers not only prevent binding of non-specific proteins, but can also be used to study the assembly of proteins of interest straight from lysate without prior purification.

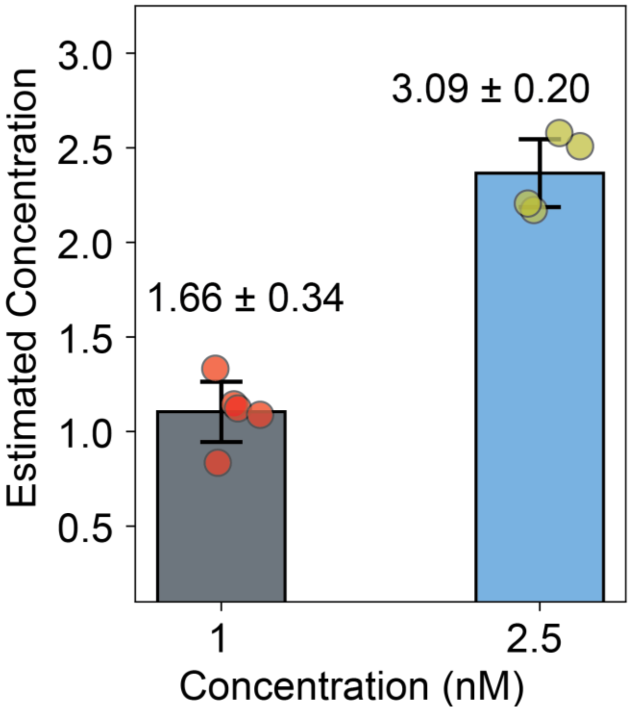


**Supplementary figure 8**. Estimation of solution concentration of purified Dyn1 using oligomeric abundance of purified Dyn1 oligomers data shown in supplementary fig. 4a.

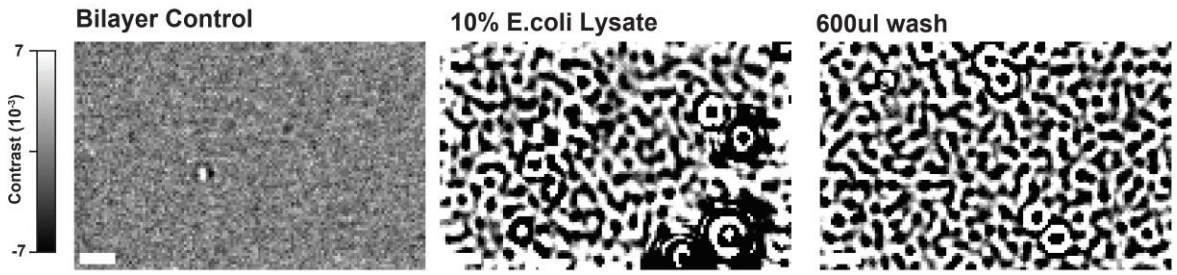


**Supplementary figure 9**. Representative background subtracted images of PS-NTA bilayers, bilayers incubated with 10% E.coli lysate and after extensively washing bilayers with reaction buffer. It is apparent that at high concentration of E.coli lysate, bilayer fail to effectively passivate against non-specific binding of E.coli proteins. Scale bar = 1µm

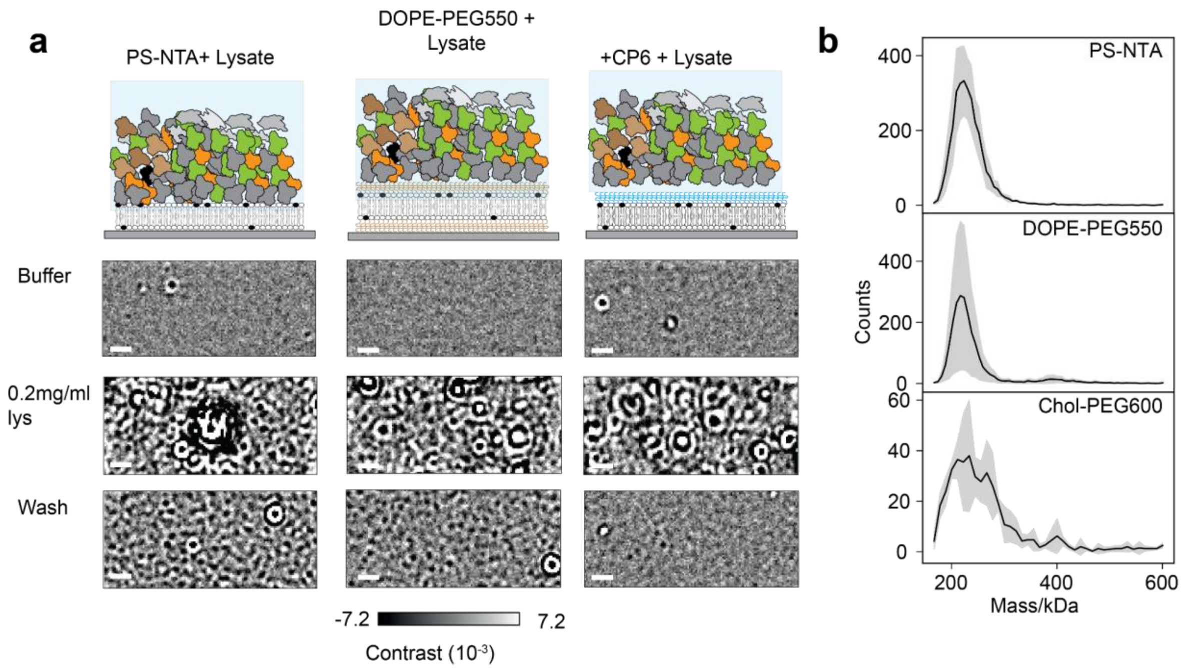


**Supplementary figure 10.** Comparison of PEG-based bilayer passivation by DOPE-PEG550 and CP6. a) Representative, background-subtracted images of PS-NTA or peg passivated PS-NTA SLBs after exposure to SHSy5y lysate. For PEG passivation comparison, PS-NTA SLBs were either supplemented with peg lipid, DOPE-PEG550 or pre-treated with CP6. 0.2mg/ml SHSy5y cell-lysate on PS-NTA SLBs (PS-NTA+ Lysate), DOPE-PEG550 SLBs (DOPE-PEG550+Lysate) and CP6 pre-treated with CP6 (+CP6+Lysate). After 5 minutes incubation, bilayers were washed with reaction buffer (wash) and imaged. Scale bar=1µM. b) Comparison of residual counts after wash. DOPE-PEG550 did not provide significant passivation over PS-NTA SLBs alone. However, CP6-based passivation, led to 10-fold lower counts of non-specifically bound proteins.

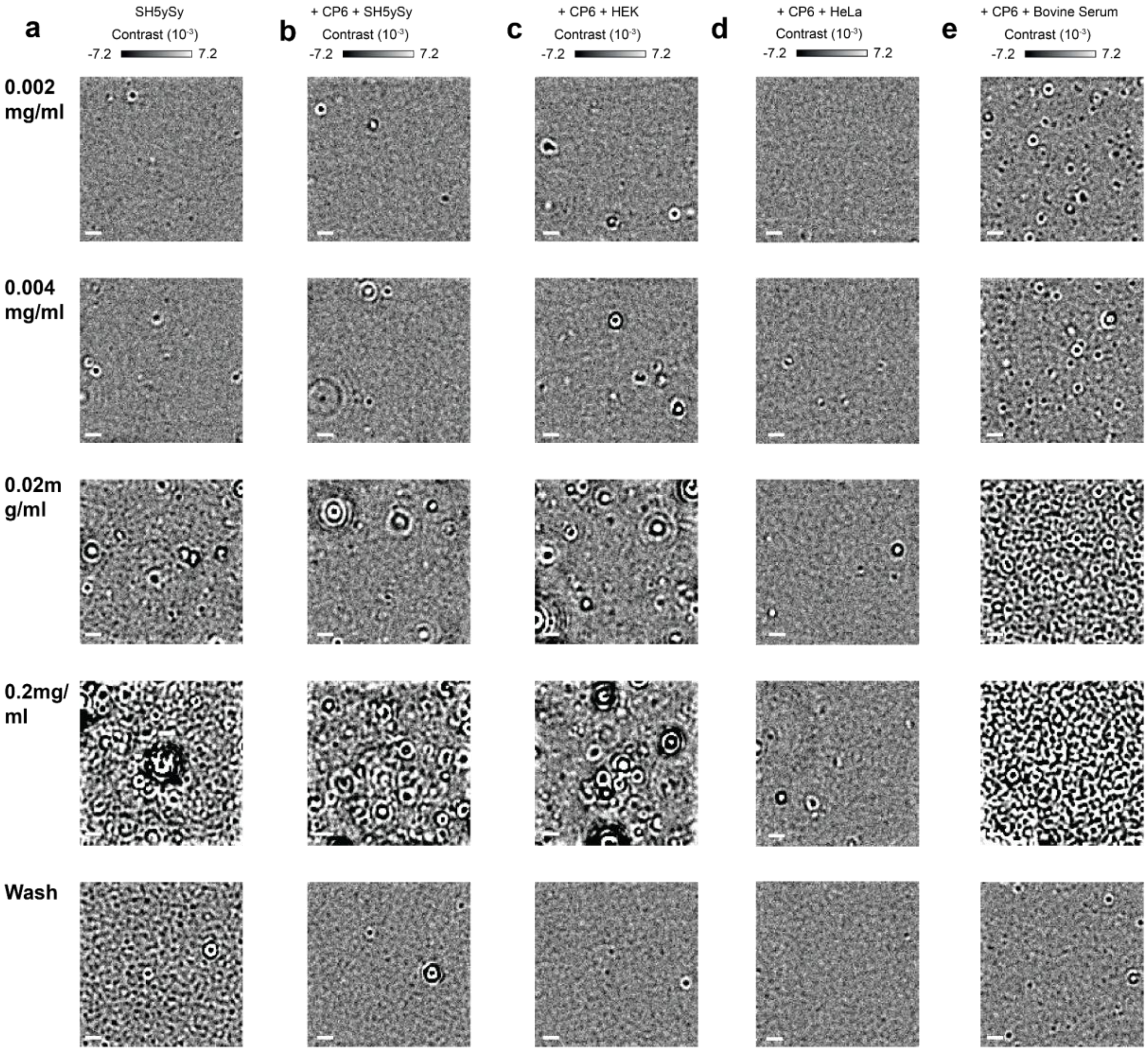


**Supplementary figure 11**. Comparison of CP6-based SLB passivation for various cell lysates. a-e) Representative background subtracted images of CP6 passivated PS-NTA SLBs after treatment with increasing cell-lysates or bovine serum concentration. Scale bar = 1µm

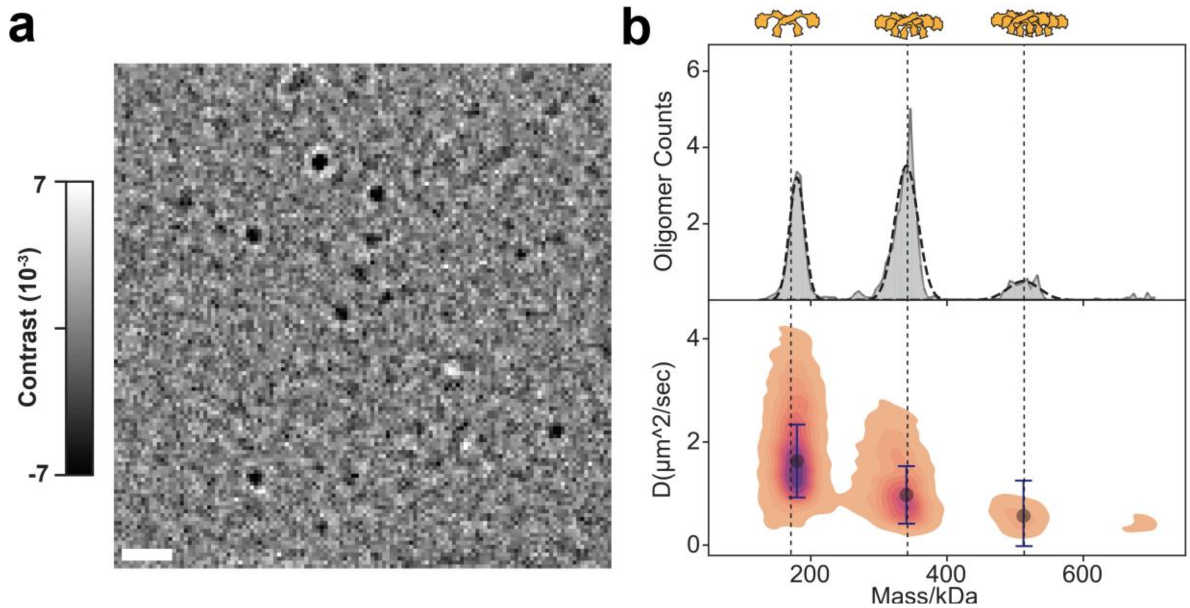


**Supplementary figure 12**. Dyn1 pulldown from E.coli lysate on CP6-fortified PS-NTA bilayers. 0.0016% E.coli lysate was incubated on bilayer and incubated for 5 minutes. a) background corrected frame showing passivated PS-NTA bilayers containing E.coli lysates and bound Dyn1 oligomers. b) Mass and diffusion plot for Dyn1 oligomers bound to passivated bilayers. 4, 1-minute movies were combined to generate mass and diffusion histogram. Shaded, vertical lines indicate the mass of Dyn1 oligomers after accounting 5 % contrast drop on bilayers. Oligomer schematic indicate the bilayer bound oligomers. Scale bar = 1 μm

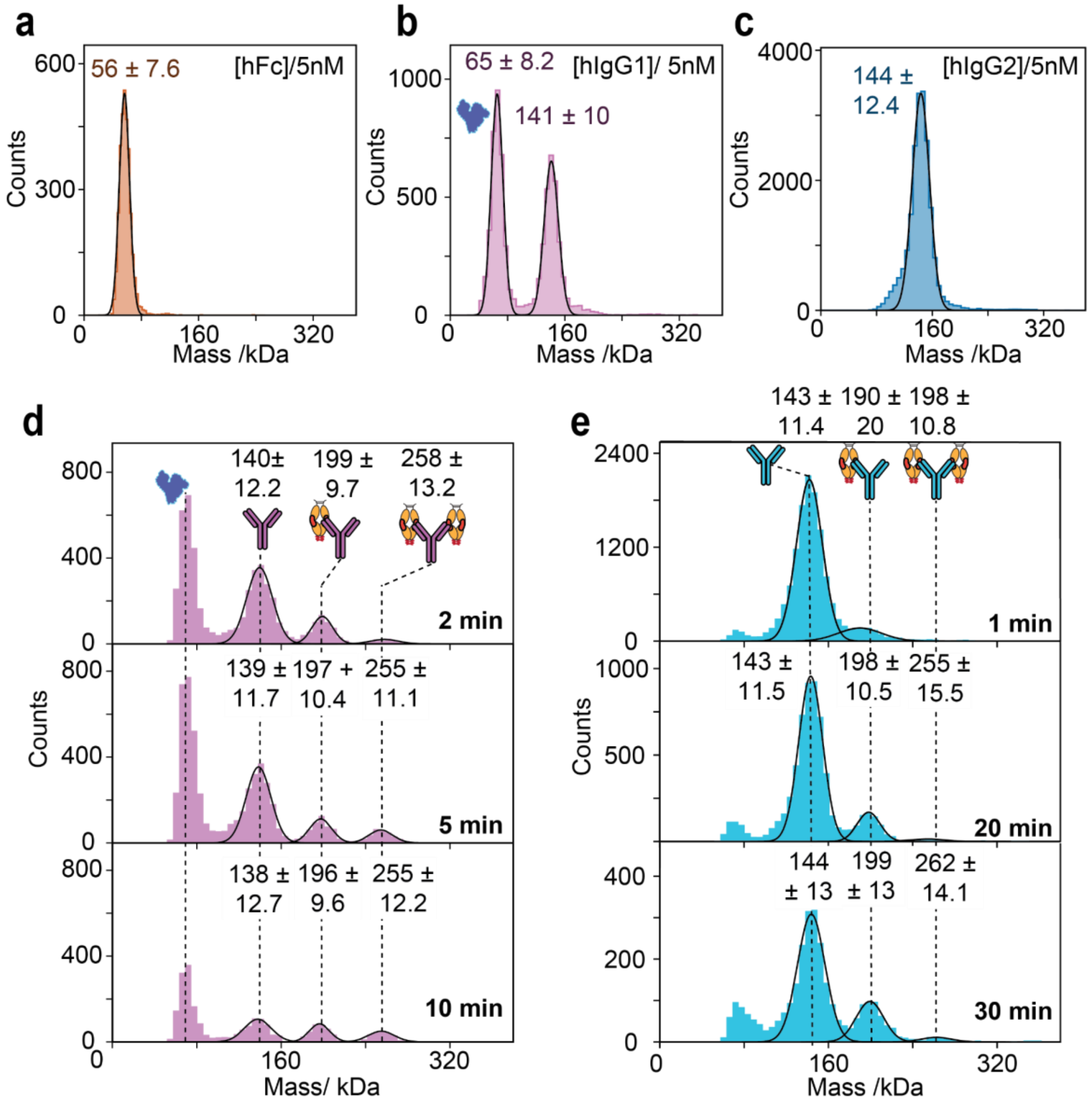


**Supplementary figure 13**. a) Mass histogram for hFc at 5 nM. Molecular weight indicates hFc is a dimer (56 kDa). However, the presence of monomer (27 kDa) can not be ruled out. b) Molecular weight distribution of hIgG1 at 5 nM. In addition to the monoclonal antibody peak at 141 kDa, sample also showed the presence of BSA (65 kDa) which was added by the manufacturer. c) Mass histogram of hIgG2 at 5 nM. Unlike hIgG1, hIgG2 did not show presence of BSA. d) Mass histogram showing hIgG1 and hFc complexes. hFc (10 nM) and hIgG1 (5 nM) were mixed and data collected at different time points. hIgG1 bound to 1 and 2 hFc dimers. e) Mass histogram showing hIgG2 (5 nM) and hFc (45 nM). Despite high hFc concentration, hIgG2 predominantly remained in free , followed by 1:1 complex of hFc:hIgG2. After 30-minute incubation at RT, we observed small fraction of 2:1, hFc:hIgG2 complex.

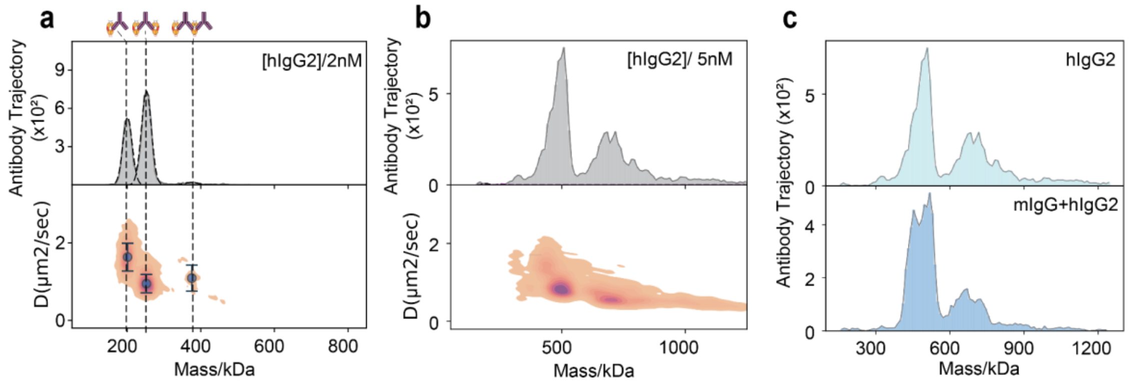


**Supplementary figure 14**. Demonstration of hIgG2 binding in the presence of non-specific Ab mIgG (100 nM ). To maximise the possibility to see non-specific binding between mIgG (100 nM) and hFc (2 nM), the interaction was carried out in the constant presence of 2nM hFc. a) Mass and diffusion distribution for hIgG2 (2 nM) without the presence of mIgG. hFc:hIgG2 complex formed show the response in the absence of mIgG (Pure). b) Mass and diffusion coefficient with 5 nM final hIgG2 concentration. Ascertaining hFc:hIgG2 complex stoichiometry from this data is complicated. c) Comparison of mass histogram of hIgG2 in the absence (hIgG2) and presence of 100 nM mIgG (mIgG+hIgG2). Comparison suggests that hIgG2 binding to hFc is not affected by the presence of non-specific mIgG at 20-fold higher concentration. The change in species is due to the presence of excess unbound hFc dimer in solution leading to complex formation both in solution and on SLBs.

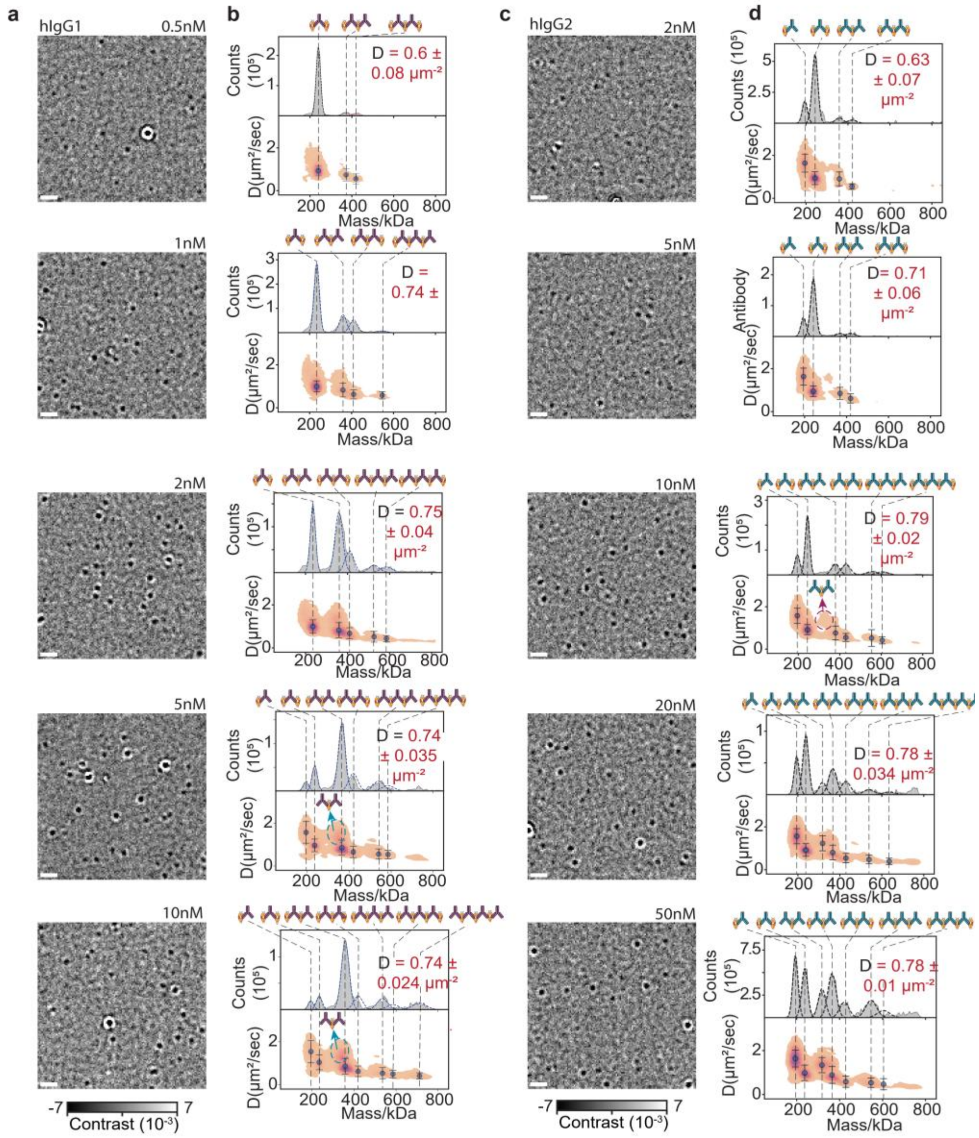

**Supplementary figure 15**. hFc, hIgG1 and hIgG2 complex stoichiometry determination. a) Titration of hIgG1 concentration added to PS-NTA slb containing hFc and CP6. Representative background subtracted images showing the complexes formed on bilayers. b) Mass and diffusion histogram showing various hFc:hIgG1 complexes formed on bilayers. for each concentration, data from 5, 1 minute movies was pooled. Mean ± SD of fitted particle density (D) from 5 movies is shown on each histogram. Molecular weight and diffusion change is used to see ascertain the stoichiometry of mass complexes, see table below. hFc:hIgG1 complex stochiometry for each species is denoted by schematic. c) Titration of hIgG2 on slbs containing hFc and CP6. Representative background subtracted images showing the complexes formed on bilayers.d) Mass and diffusion histogram showing various hFc:hIgG2 complex formation as a function of increasing hIgG2 concentration. For each hIgG concentration, data is pooed from 5, 1-minute movies. D represents the mean ± SD of fitted particle density from 5, 1-minute movies.

**a**

| Species | Mass/kDa (Mean ± SD) | D1/μm².sec⁻¹ (Mean ± SD) |
|---|---|---|
| 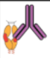 | 194 ± 0.5 | 1.5 ± 0.5 |
| 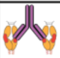 | 235.7 ± 2.4 | 1 ± 0.1 |
| 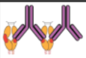 | 362.6 ± 6.2 | 0.82 ± 0.09 |
| 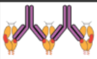 | 415.4 ± 5.4 | 0.64 ± 0.05 |
| 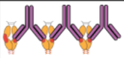 | 537 ± 8.6 | 0.55 ± 0.05 |
| 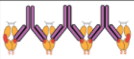 | 585.3 ± 9 | 0.47 ± 0.05 |
| 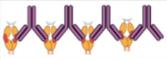 | 712 ± 10 | 0.4 ± 0.05 |

**b**

| Species | Mass/kDa (Mean ± SD) | D1/μm².sec⁻¹ (Mean ± SD) |
|---|---|---|
| 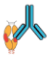 | 194.3 ± 0.3 | 1.6 ± 0.03 |
| 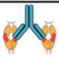 | 240.5 ± 2.4 | 0.9 ± 0.042 |
| 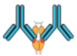 | 320.4 ± 3.7 | 1.3 ± 0.07 |
| 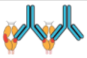 | 367.9 ± 6.9 | 0.84 ± 0.05 |
| 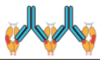 | 422.8 ± 9.3 | 0.58 ± 0.04 |
| 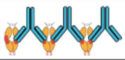 | 546.2 ± 7.3 | 0.52 ± 0.03 |
| 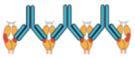 | 606.2 ± 3.2 | 0.43 ± 0.06 |

**Supplementary figure 16**. Molecular weight and diffusion relation of various hFc:hIgG1 and hIgG2 complexes. For mass and diffusion, mean ± SD of fitted mass and diffusion coefficient is plotted.

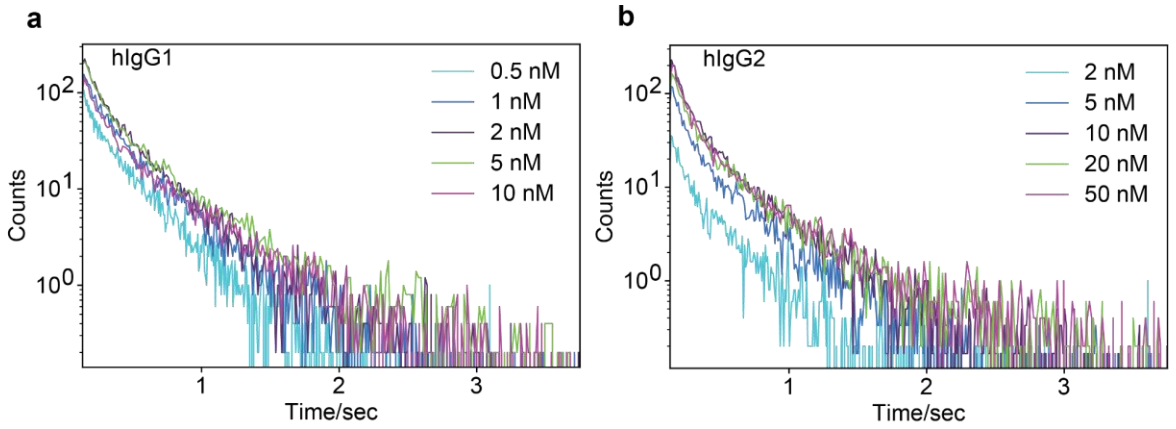


**Supplementary figure 17**. Trajectory length distribution for data shown in supplementary figure 14. For hIgG1/2, trajectory length for all particles between 0 and 1250 kDa for each concentration was pooled and plotted. Distribution of trajectory length shows that with increasing hIgG concentration, the trajectory length remains unchanged.

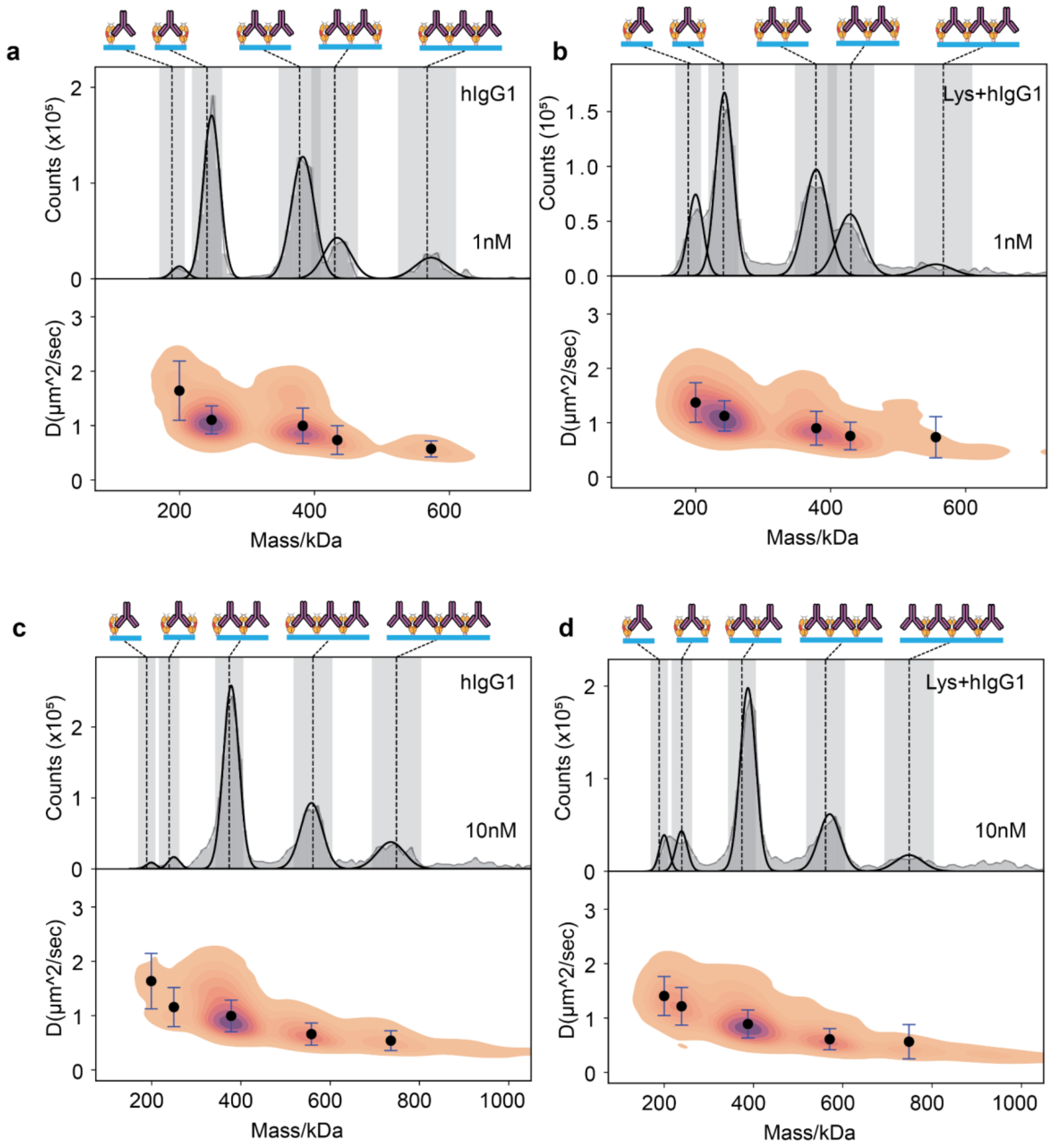


**Supplementary figure 18**. a-b) Mass and diffusion distribution for hFc-hIgG1 complex formation when 1 nM hIgG1 is incubated in the absence (a) and presence of lysate (b) with CP6 passivated PS-NTA SLBs for 5 minutes, washed and measured. c-d) Mass and diffusion distribution for hFc-hIgG1 complex formation when 10 nM hIgG1 is incubated in the absence (a) and presence of lysate (b) with CP6 passivated PS-NTA SLBs for 5 minutes, washed and measured. Data from 4, 1 minute movies for each condition was pooled. In case of 10 nM hIgG1, large complexes where increasing number of hFc was crosslinked by hIgG1. The abundance of crosslinked hFc matched between purifed hIgG1 and hIgG1 mixed with lysate.

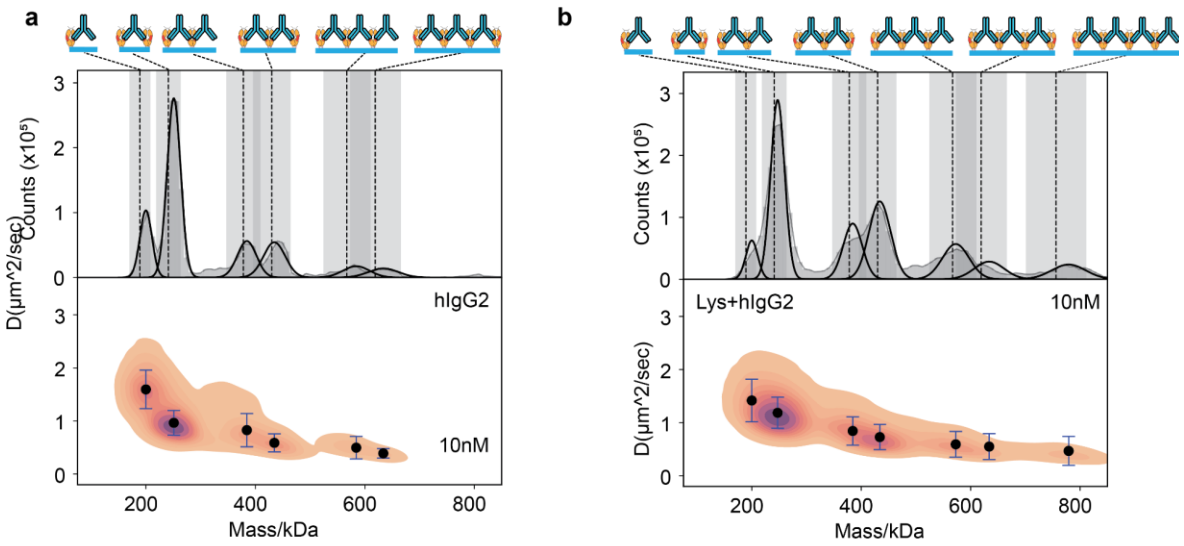


**Supplementary figure 19**. a-b) Mass and diffusion distribution for hFc-hIgG2 complex formation when 1 nM hIgG1 is incubated in the absence (a) and presence of lysate (b) with CP6 passivated PS-NTA SLBs for 5 minutes, washed and measured. Data from 6, 1 minute movies for each condition was pooled. Like hIgG1, hIgG2 showed similar complex abundance between pure hIgG2 (a) and hIgG2 mixed with lysate(b). In contrast to 10nM hIgG1, hIgG2, however, showed equal abundance of 2&3 and 3&4 hFc crosslinked by hIgG2.

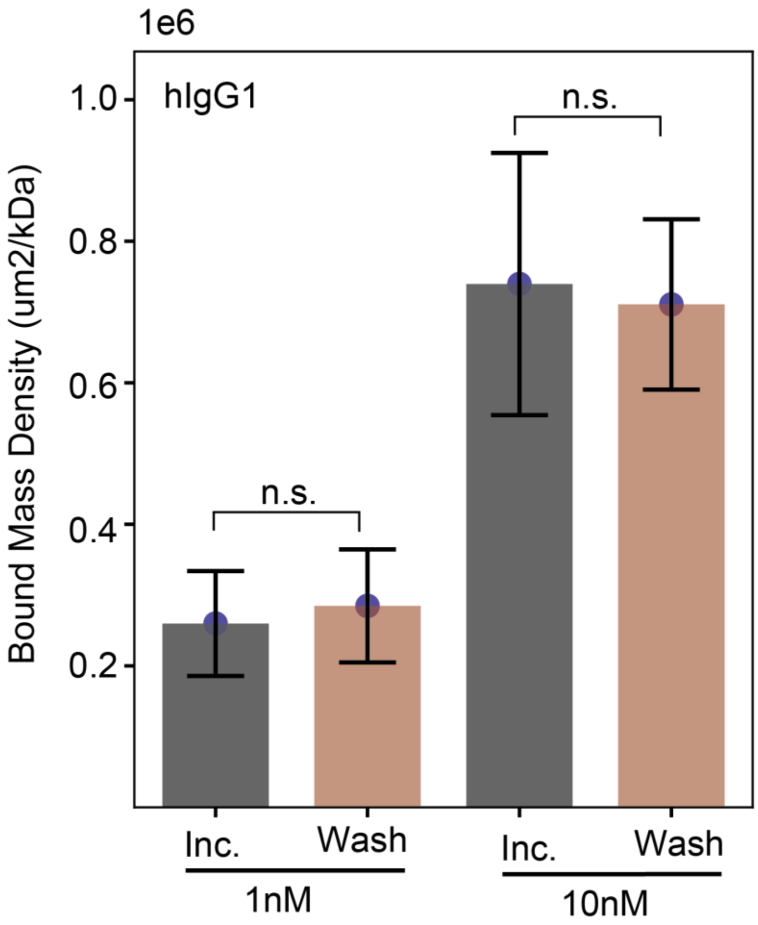


**Supplementary figure 20**. a-b) Mass and diffusion distribution for hIgG1-hFc complex formation in the absence (a) and presence of lysate (b). 1 nM hIgG1 was incubated with or without lysate, on CP6 fortified PS-NTA SLBs for 5 minutes, washed and measured. Data from 4, 1-minute movies for each condition was pooled.
